# In pursuit of a better broiler: A comparison of the inactivity, behavior, and enrichment use of fast- and slower-growing broiler chickens

**DOI:** 10.1101/2021.05.18.444545

**Authors:** Lauren C. Dawson, Tina M. Widowski, Zhenzhen Liu, A. Michelle Edwards, Stephanie Torrey

**Affiliations:** Department of Animal Biosciences, University of Guelph, Guelph ON, Canada; Campbell Centre for the Study of Animal Welfare, University of Guelph, Guelph ON, Canada; Ontario Agricultural College, University of Guelph, Guelph ON, Canada

**Keywords:** Broiler, Welfare, Inactivity, Behavior, Slow-growing

## Abstract

Selection for rapid growth has produced heavier, more efficient broiler chickens, but has also introduced health and welfare issues, which may cause or be caused by inactivity. Rapid growth may also limit the performance of motivated behaviors, whereas the provision of enrichment may increase these behaviors and general activity. This study aimed to evaluate the inactivity, behavior, and enrichment use of 2 fast- (**CONV**) and 12 slower-growing broiler strains [categorized as fastest (**FAST**), moderate (**MOD**) and slowest slow (**SLOW**), based on their growth rates]. To evaluate inactivity, one male and one female from 153 pens were outfitted with omni-directional accelerometers from day 21 until processing (14–24 birds/strain from 8-12 pens/strain). Additionally, to supplement inactivity data, five-minute continuous behavioral observations of four focal birds per pen (2 males, 2 females) were conducted on days 26, 42, and 56 (72–148 observations of 8–12 pens/strain) to quantify the duration and frequency of various behaviors; at the same time, 5–11 instantaneous scan samples were also performed for pen-based enrichment use. Inactivity peaked at 78-80% of the day for all strains; however, those with slower growth rates reached these levels at older ages. Compared to slower-growing strains at the same age, faster-growing strains were more inactive, spent more time sitting and feeding, spent less time standing and walking, and used enrichments less; these differences mostly occurred at younger ages. Generally, at the same age, strains with similar growth rates (within the same category) behaved similarly, with only a few exceptions. Results suggest that not all strains identified as ‘slow-growing’ broilers behave differently from fast-growing broilers, nor do they all behave similarly to each other. As such, results suggest that improved broiler welfare, particularly with respect to reduced inactivity, the performance of a wider range of normal, motivated behaviors, and/or increased enrichment use, is related to the broiler strain’s specific growth rate.

## INTRODUCTION

Due to genetic selection for rapid growth and advances in management practices, modern broiler chickens now reach market weight (> 2 kg) in as little as 6 weeks. In comparison to broiler chickens from the 1950s, current commercial broilers have higher body weights, increased breast yields, and improved feed conversion ratios (Havenstein et al., 2003a; b; Zuidhof et al., 2014). These improvements in production and efficiency, however, come at a cost to broiler health and welfare. Modern broilers have poorer immune system function (Cheema et al., 2003) and higher mortality rates (Havenstein et al., 2003a). Rapid growth is also associated with increased incidences of metabolic disorders (Bessei, 2006), muscle myopathies [e.g. white striping, woody breast (Kuttappan et al., 2016)], skeletal disorders [e.g. varus/valgus deformities (Shim et al., 2012), tibial dyschondroplasia (**TD**; Fanatico et al., 2008; Shim et al., 2012)], and lameness (Kestin et al., 2001; Wilhelmsson et al., 2019).

Fast-growing (‘conventional’) broilers also have low activity levels. Broilers can spend as much as 70–80% of their day sitting (Bizeray et al., 2000; Weeks et al., 2000; Bokkers and Koene, 2003; Dixon, 2020), whereas their ancestors, red junglefowl, spend as little as 10% of their day sitting (Dawkins, 1989). Low locomotor activity, or inactivity, is not a welfare concern in and of itself; however, inactivity may cause or be caused by broiler welfare issues. For one, low activity is linked to poor leg health and lameness. Physical exercise improves bone development and health, and reduces deformities (Reiter and Bessei, 1998, 2011), whereas a lack of exercise may lead to a higher incidence of leg deformities (Haye and Simons, 1978). Based on lame broilers’ behavioral responses to analgesics (McGeown et al., 1999; Danbury et al., 2000), lameness is painful, so broilers who become lame due to a lack of exercise are likely to further decrease their activity. Low activity is also linked to contact dermatitis. Low locomotor activity implies an increased duration of time sitting or lying in contact with potentially poor quality litter, which increases the incidence of contact dermatitis (i.e. foot-pad dermatitis, hock burn; Bassler et al., 2013; De Jong et al., 2014). The lesions associated with contact dermatitis are also thought to be painful (Bessei, 2006; Haslam et al., 2006).

Beyond general inactivity, the specific behaviors that broilers perform, in terms of both the duration of time spent performing them and the frequency with which they are performed, may also be altered by their growth rate. For example, in addition to differences in sitting, fast- and slower-growing broilers also differ in the time they spend standing, walking, feeding, drinking, and performing exploratory behavior (Bizeray et al., 2000; Bokkers and Koene, 2003; Wallenbeck et al., 2016; Dixon, 2020). Broilers’ heavy body weight and/or conformation may also prevent their engagement in normal behaviors, or prevent the performance of other behaviors that broilers are motivated to perform, such as perching or dustbathing (Bokkers and Koene, 2003; Wallenbeck et al., 2016; Dixon, 2020). Altogether, if broilers are unable to perform normal, highly-motivated behaviors, they may become frustrated (Bradshaw et al., 2002; Bokkers et al., 2007).

The provision of environmental enrichment may improve broiler welfare by increasing activity, while also providing an opportunity for the expression of a wider range of normal, motivated behaviors (e.g. Bizeray et al., 2002; Bach et al., 2019; Vasdal et al., 2019). Elevated platforms allow for elevated resting, a natural behavior for junglefowl, and their provision has been linked to improved walking ability (i.e. better gait scores; Kaukonen et al., 2017), and lower incidences of TD (Kaukonen et al., 2017) and footpad dermatitis (Tahamtani et al., 2020). Elevated platforms may also increase useable space while reducing time in contact with the litter (Bizeray et al., 2002), improve thermoregulation (Riber et al., 2018), reduce fearfulness (Tahamtani et al., 2018), and increase the performance of comfort behaviors (i.e. dustbathing, preening, stretching; Bach et al., 2019). Additionally, oral enrichment objects (e.g. hanging strings, straw bales) increase the opportunity for pecking and foraging, and have also been linked to improved gait scores (Bailie and O’Connell, 2015) and longer latencies to lie (Bailie et al., 2013).

Given the welfare issues associated with rapid growth, there has been increasing interest in the potential of using slower-growing broiler strains for commercial production. Although previous studies have evaluated the welfare or welfare-related outcomes of fast- and slower-growing broilers (e.g. Bokkers and Koene, 2003; Fanatico et al., 2005; Lichovníková et al., 2017; Wilhelmsson et al., 2019; Dixon, 2020; Mancinelli et al., 2020; Weimer et al., 2020), many of these studies directly compared a limited number of strains (e.g. 1 conventional vs. 1 slower-growing strain: Bokkers and Koene, 2003; Wallenbeck et al., 2016; Lichovníková et al., 2017; Wilhelmsson et al., 2019; Weimer et al., 2020), evaluated welfare in non-intensive settings (e.g. Fanatico et al., 2005; Mancinelli et al., 2020), and/or evaluated slower-growing birds at comparably low body weights (e.g. < 2 kg: Bokkers and Koene, 2003; Lichovníková et al., 2017). Additionally, even among strains considered to be slower growing, there is a large range in growth rates and no consensus about growth rate cut-offs that ensure or lead to improved welfare.

Our goal was to benchmark the welfare of broiler chickens with different growth rates, including various degrees of slow growth, raised to similar body weights under the same management and housing conditions. To do so, we developed a large scale, multi-outcome study designed to evaluate the effects of both age and body weight on the welfare outcomes of 2 conventional and 12 slower-growing broiler strains. Differences in efficiency and mortality are available in Torrey et al. (2021). This paper presents the results of a sub-section of the larger study, with a focus on behavioral measures of welfare. More specifically, across 14 broiler strains, we monitored and compared inactivity levels using accelerometers, assessed differences in the durations and frequencies of behaviors via live observations, and compared enrichment use using scan sampling.

## MATERIALS AND METHODS

### Animals, Management and Housing

Animal use was approved by the University of Guelph Animal Care Committee (Animal Utilisation Protocol #3746). This study encompasses a subsection of a large-scale study to investigate behavioral, physiological, and production differences between conventional (fast-growing) and slower-growing broiler strains. A full description of the animals, management and housing is available in Torrey et al. (2021).

In short, the study followed a randomized incomplete block design, with 8 trials and up to 28 pens per trial. All pens were in the same room at the Arkell Poultry Research Facility (Guelph, ON, Canada), and this room was used for all trials. Pens were divided into four blocks according to location to account for differences in microclimate within the room. In total, 14 broiler strains were tested: 2 conventional and 12 slower-growing strains. Each strain was assessed in up to 3 trials, in 4 pens per trial, and in 1 pen per block per trial.

All strains were raised under similar conditions: in a 238 × 160 cm pen with softwood shavings as litter, 5 nipple drinkers, and a hanging feeder with *ad libitum* feed formulated for slow growth. Pens were enriched with a 30-cm high platform with a 25° ramp, a hanging scale, ¼ of a PECKstone mineral block (Protekta, Lucknow, Ontario, Canada), and a hanging nylon rope with strips of polyester cloth tied to the end. Broilers were placed in pens of 44 birds (22 females, 22 males) on day of hatch. Group sizes were reduced to 38 birds to maintain a stocking density of 30 kg/m^2^. Light intensity was kept at 20 lux. To assess the impact of both age and weight, strains were processed at different ages, according to breeders’ expected time to reach one of two target weights (**TW**): 2.1 kg (TW 1) or 3.2 kg (TW 2). Conventional strains were processed at day 34 (TW 1) or day 48 (TW 2) and slower-growing strains were processed at day 48 (TW 1) or day 62 (TW 2).

### Inactivity

Actical^®^ (Philips Respironics, Murrysville, PA, USA) accelerometers were used to continuously assess broiler activity. Actical^®^ accelerometers are small (28 × 27 × 10 mm) and lightweight (17.5 g), with a sensitivity of 0.05–2 G and frequency of 0.35–3.5 Hz. They are multidirectional piezoelectric sensor accelerometers that can detect movement along any axis, accounting for both the intensity and duration of movement. These specific accelerometers have been used to assess activity in a number of species, including laying hens (Casey-Trott and Widowski, 2018), dairy cattle (Rialland et al., 2014), cats (Lascelles et al., 2008; de Godoy and Shoveller, 2017), dogs (Mejia et al., 2019; Phungviwatnikul et al., 2020), and rhesus monkeys (Papailiou et al., 2008), and have been validated for assessing inactivity in laying hens (Casey-Trott and Widowski, 2018). Acticals^®^ were programmed as follows: epoch length: 1 s; subject height: 10.0 cm; subject weight 0.5 kg; subject sex: female; subject age: 2; these settings maximize sensitivity for activity detection in laying hens (Casey-Trott and Widowski, 2018).

At 21 days of age, one male and one female bird per pen were outfitted with an Actical^®^ accelerometer; at this age, the device weighed less than 5% of the body weight of birds from all strains, per the recommendation of Siegford et al. (2016) for wearable equipment. Moreover, weight-matched broilers with and without Actical^®^ devices installed behave similarly during mobility tests, suggesting that their use does not affect broiler behavior (Liu, 2019). Birds were randomly selected from 12 sentinel birds (that were selected at hatch) and were not subject to any other live testing, other than behavioral observations. Acticals^®^ were attached as ‘backpacks’ using beige-colored cotton elastic material that was looped through the device and around the birds’ wings, with two fingers’ worth of space between the strap and wing. Straps were checked every few days and adjusted as needed to ensure that they did not affect circulation or movement. Chickens wore the accelerometers continuously until the day prior to processing.

Activity data was downloaded from the Actical^®^ devices using Actical 3.10 software (Koninklijke Philips N.V., Amsterdam, The Netherlands). Using SAS 9.4, the number of 15-second epochs with no activity counts (zero acceleration registered) were summed per day to calculate daily inactivity counts for each bird. The day of installation and removal were excluded from analysis to avoid including incomplete data.

### Behavioral Observations

At 26, 42, and 56 days of age, behavioral observations were conducted on four focal birds per pen (two males, two females, including the two birds outfitted with an Actical^®^ accelerometer, all selected from the 12 sentinel birds selected at hatch). Observations were made according to the ethogram outlined in Supplementary Table 1 (see Table 1 for a simplified version). All occurrences of each state and event behavior, as well as the location in which they occurred, were recorded using The Pocket Observer v3.2.43 (Noldus Information Technology, Wageningen, The Netherlands) on a Samsung Galaxy Tab4 tablet (Seoul, South Korea). Each focal bird was observed continuously for 5 minutes, and all four birds per pen were successively and systematically observed to reduce bias towards activity. Observations took place between 08h00 and 14h30. All observers (n = 10 total across the different trials) were trained and had acceptable inter-observer reliability (kappa > 0.60; McHugh, 2012). When multiple observers were used on the same day, each observer watched all pens within a block(s) to ensure that no single observer assessed all the pens of one strain.

**Table 1.**
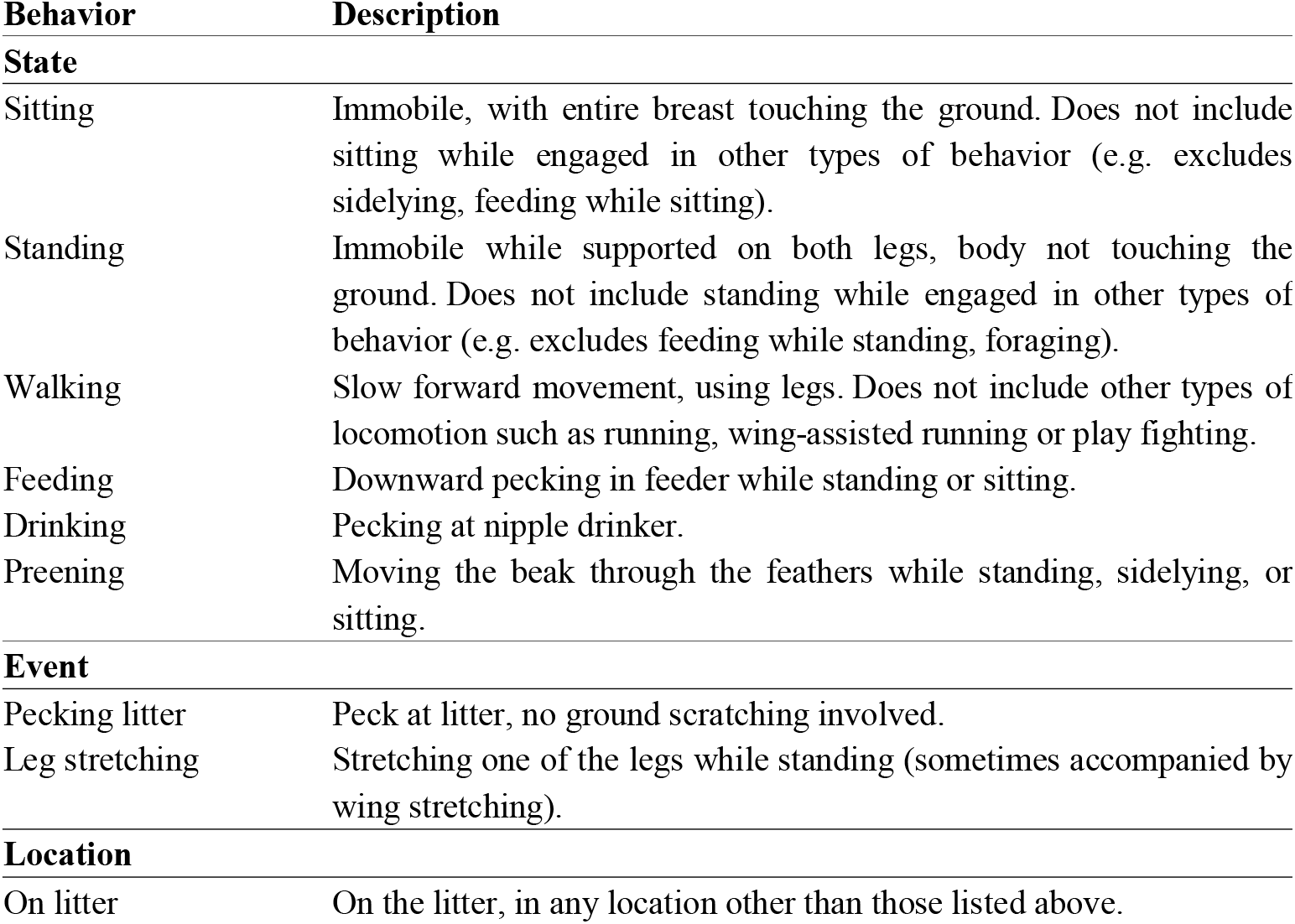
Simplified ethogram for live behavioral observations.

| **Behavior** | **Description** |
| --- | --- |
| **State** |  |
| Sitting | Immobile, with entire breast touching the ground. Does not include sitting while engaged in other types of behavior (e.g. excludes sidelying, feeding while sitting). |
| Standing | Immobile while supported on both legs, body not touching the ground. Does not include standing while engaged in other types of behavior (e.g. excludes feeding while standing, foraging). |
| Walking | Slow forward movement, using legs. Does not include other types of locomotion such as running, wing-assisted running or play fighting. |
| Feeding | Downward pecking in feeder while standing or sitting. |
| Drinking | Pecking at nipple drinker. |
| Preening | Moving the beak through the feathers while standing, sidelying, or sitting. |
| **Event** |  |
| Pecking litter | Peck at litter, no ground scratching involved. |
| Leg stretching | Stretching one of the legs while standing (sometimes accompanied by wing stretching). |
| **Location** |  |
| On litter | On the litter, in any location other than those listed above. |

The total duration spent performing each state behavior and the time spent in each location, as proportions of the total observation time, the number of bouts of each state behavior and the sum frequency of each event behavior were compiled. Many behaviors were performed in a small number of observations, and only behaviors that occurred in ≥ 20% of all observations were analyzed statistically; these behaviors were: sitting, standing, walking, feeding (feed standing + feed sitting), drinking, and preening (preen standing + preen sidelying + preen sitting). With respect to location, on the litter (on litter + under platform/ramp) was the only location that occurred frequently enough to be analyzed; it was assessed in terms of duration only. Moreover, the total duration spent sitting in any capacity (‘all sitting’ = sitting + feed sitting + preen sitting), standing in any capacity (‘all standing’ = standing + feed standing + preen standing + drinking), in locomotion (walking + running + wing-assisted running), engaging in any behavior in which the chicken was off their feet (‘off feet’ = all sitting + sidelying + preen sidelying + dustbathing) and on their feet (‘on feet’ = all standing + locomotion + playfighting + foraging) were also calculated for analysis.

### Enrichment Use

Prior to the behavioral observation of each pen (at ∼20–25 min intervals), an observer performed an instantaneous scan of their block(s) of observation pens to assess the number of birds on the platform, on the ramp, under the platform or ramp, on the scale, on the pecking stone, pecking the pecking stone, and pecking the rope. Locations were assigned based on the location of the chicken’s two feet (e.g. both feet on the ramp = on the ramp) and/or the location of the majority of the body (e.g. > 50% under the platform = under the platform). A minimum of five scans were performed of each pen on each observation day, by the same observer who had conducted the behavioral observations for that pen. Since scans were performed prior to each observation, the number of scans performed on any given day varied with the number of occupied pens in the room and the number of observers performing observations on that day; however, across the whole study, the mean number of scans was approximately the same for all strains. On a pen basis, the proportion of birds engaged in the use of each enrichment was calculated. Additionally, the proportion of birds using any enrichment (sum of all enrichment use), as well as the proportion of birds on the litter (total number of birds – number of birds using any enrichment + number of birds under the platform or ramp) were calculated. As was the case for behavioral observations, some enrichment use (i.e. on the pecking stone, pecking the pecking stone, pecking the rope) occurred infrequently (< 5% of all scans) and was not analyzed statistically.

### Statistical Analysis

To facilitate analyses, the 14 strains were categorized into four groups according to ADG to TW 2, as follows: conventional (**CONV**; strains B and C; ADG_0-47_ = 66.0-68.7 g/day), fastest slow (**FAST**; strains F, G, I, M; ADG_0-61_ = 53.5-55.5 g/day), moderate slow (**MOD**; strains E, H, O, S; ADG_0-61_ = 50.2-51.2 g/day), and slowest slow (**SLOW**; strains D, J, K, N; ADG_0-61_ = 43.6-47.7 g/day). Results from each of the three sets of data (i.e. inactivity, behavioral time budgets and frequencies, and enrichment use) were compared between categories, to assess differences at different growth rates, and between strains within categories, to assess differences at similar growth rates. Comparisons were made at the same age and at the same target weight, and models to evaluate the effects of age and body weight on each outcome were run separately.

Inactivity was analyzed using generalized linear mixed models in SAS 9.4, with pen as the experimental unit, and block and block nested within trial as random effects. CONV birds were only raised until week 7, producing holes in the data for weeks 8 and 9; this necessitated running two models to evaluate the effect of age across categories: weeks 4 to 7, for all categories; and weeks 8 and 9, for slower-growing categories only. Four additional age models (one model per category) were built to compare differences within the same category over time, to assess when inactivity ceased to continue to increase; again, this was necessary due to the holes in the data. All age models used daily inactivity counts and week of age as a repeated measure, with a compound symmetry covariance structure, as it produced the lowest Aikaike information criterion value. The target weight model used the mean daily inactivity count from the 7 days prior to processing. For all inactivity models, fixed effects were category, strain nested within category, and bird sex; target weight or week of age was also included as a fixed effect, depending on the model in question. Interactions included in both the age and weight models were sex×category and sex×strain(category). The age models also included age×category and age×strain(category), whereas the weight model also included target weight×category and target weight×strain(category).

Behavior and enrichment use were analyzed using generalized linear mixed models in SAS 9.4, with pen as the experimental unit, and block and block nested within trial as random effects. Again, since CONV birds were raised until day 48, there were holes in the data for the observations performed on day 56. As such, two models were built to evaluate the effect of age: days 26 and 42, for all categories, with age as a repeated measure and using the covariance structure that produced the lowest Aikaike information criterion value; and day 56, for slower-growing categories only. For all behavioral observation models, the fixed effects were identical to those listed for the inactivity models, except for the addition of observer as a fixed effect. The fixed effects were also similar in the enrichment use models, with the addition of observer and the removal of bird sex and its interactions, since observations were conducted on a pen basis, with pens including both male and female birds.

For all models, linearity and the homogeneity of effects were assessed using scatterplots and boxplots of studentized residuals. Normality was assessed using quantile-quantile plots and Shapiro-Wilk tests. Selection of the most appropriate distribution was based on residual analysis; thus, the selected distribution varied depending on the model so as to meet model assumptions. The Kenward Roger method was used to compute denominator degrees of freedom. P-values were adjusted for multiple comparisons using the Tukey adjustment. Significance was set at an adjusted P-value < 0.05. To facilitate the interpretation of inactivity results, back-transformed least-square mean inactivity counts were converted into minutes, wherein 1 inactivity count equals 15 seconds of inactivity.

## RESULTS

### Inactivity

Data from 153 pens, accounting for 138 males and 142 females, were analyzed. CONV were represented in 20 pens, FAST in 42 pens, MOD in 47 pens, and SLOW in 44 pens. Each strain was represented by 14–24 birds in 8-12 pens.

#### By Age

Inactivity increased as the birds aged. As shown in Figure 1, inactivity reached a maximum of 1134 minutes, or 78% of the day, at week 7 for conventional strains and approximately 1150 minutes, or 80% of the day, at week 9 for slower-growing strains. Although all categories eventually reached similar levels of inactivity, the age at which inactivity ceased to increase differed by category. Week to week, CONV inactivity levels increased until week 6 (all pairwise P < 0.01 for weeks 4 to 6), after which point inactivity did not differ (week 6 vs. 7: P = 0.8437). Similarly, FAST inactivity increased until week 8 (all pairwise P < 0.0001 for weeks 4-8), MOD inactivity increased until week 6 (all pairwise P < 0.01 for weeks 4-6), and SLOW inactivity increased until week 8 (all pairwise P < 0.0001 for weeks 4-8).

**Figure 1.**
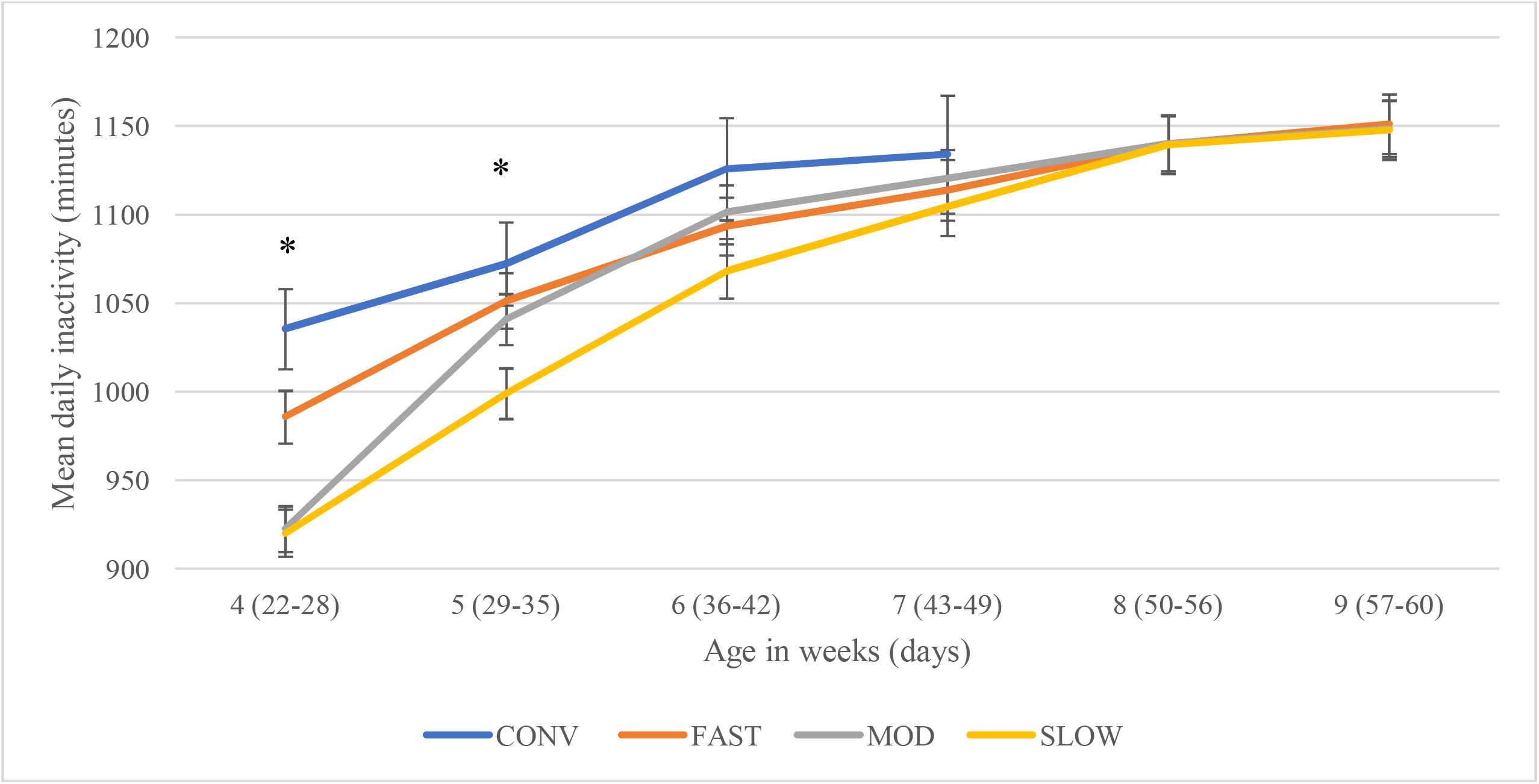
Mean daily inactivity levels (back-transformed LS-means ± SEM) by category at the same age. Broiler chickens within CONV (2 strains; ADG_0-47_ = 66.0-68.7 g/day), FAST (4 strains; ADG_0-61_ = 53.5-55.5 g/day), MOD (4 strains; ADG_0-61_ = 50.2-51.2 g/day), and SLOW (4 strains; ADG_0-61_ = 43.6-47.7 g/day) strains were compared at the same age, as measured using an omni-directional accelerometer (Actical^®^). * At the same age, categories differ at adjusted P < 0.05.

Differences between categories were greatest at younger ages (Figure 1). At week 4, CONV were more inactive than MOD (P < 0.0001) and SLOW (P < 0.0001), equating to differences of 113 and 115 minutes per day, respectively. At week 4, FAST were more inactive than MOD (P = 0.0068) and SLOW (P = 0.0054), with differences of 63 and 66 minutes per day, respectively. At week 5, CONV were more inactive than SLOW (P = 0.0410); CONV were inactive for 1072 ± 23 minutes per day, whereas SLOW were inactive for 999 ± 14 minutes per day, or 73 minutes per day less than CONV. Starting from week 6, all categories had similar levels of inactivity.

Within categories, MOD strains had different levels of inactivity, but only at younger ages. At week 4, one strain (H) had lower daily inactivity levels than the three other MOD strains (E, O and S; all pairwise P < 0.0001); H was inactive for 710 ± 21 minutes per day, whereas the mean inactivity level of E, O and S was 1007 minutes per day, equating to a difference of 296 minutes. At week 5, H tended to be less inactive than E and S (P = 0.0962 and 0.0602, respectively); H was inactive for 128 minutes less than the mean daily inactivity level of E and S. At all other ages, MOD strains did not differ. Conversely, within each category and at the same age, CONV, FAST, and SLOW strains had similar levels of inactivity; this was true for all ages.

Overall, females were more inactive than males; this amounted to a difference of 203 minutes per day during weeks 4-7, and 139 minutes per day during weeks 8–9 (P < 0.0001 for both). There were significant interactions between sex and category, as well as between sex and strain nested within category (see Supplementary Material and Supplementary Tables 2–3).

#### By Target Weight

Inactivity tended to increase as body weight increased (F_1,14_ = 3.16, P = 0.0977). CONV birds were inactive for 96 more minutes at TW 2 than at TW 1 (P = 0.0447). For FAST, MOD, and SLOW birds compared across target weights, activity levels at TW1 did not differ from those at TW2 (all P > 0.05).

Differences between categories only existed at the lighter target weight (Figure 2). At TW 1, CONV had a mean daily inactivity level of 1028 ± 22 minutes and were less inactive than FAST (P = 0.0002), MOD (P < 0.0001), and SLOW (P = 0.0021); these differences corresponded to 79, 98 and 64 minutes, respectively. At TW 2, all categories had similar mean daily levels of inactivity, ranging from 1124 to 1145 minutes per day, or 78–79% of the day. Within categories, at the same weight, strains had similar levels of inactivity.

**Figure 2.**
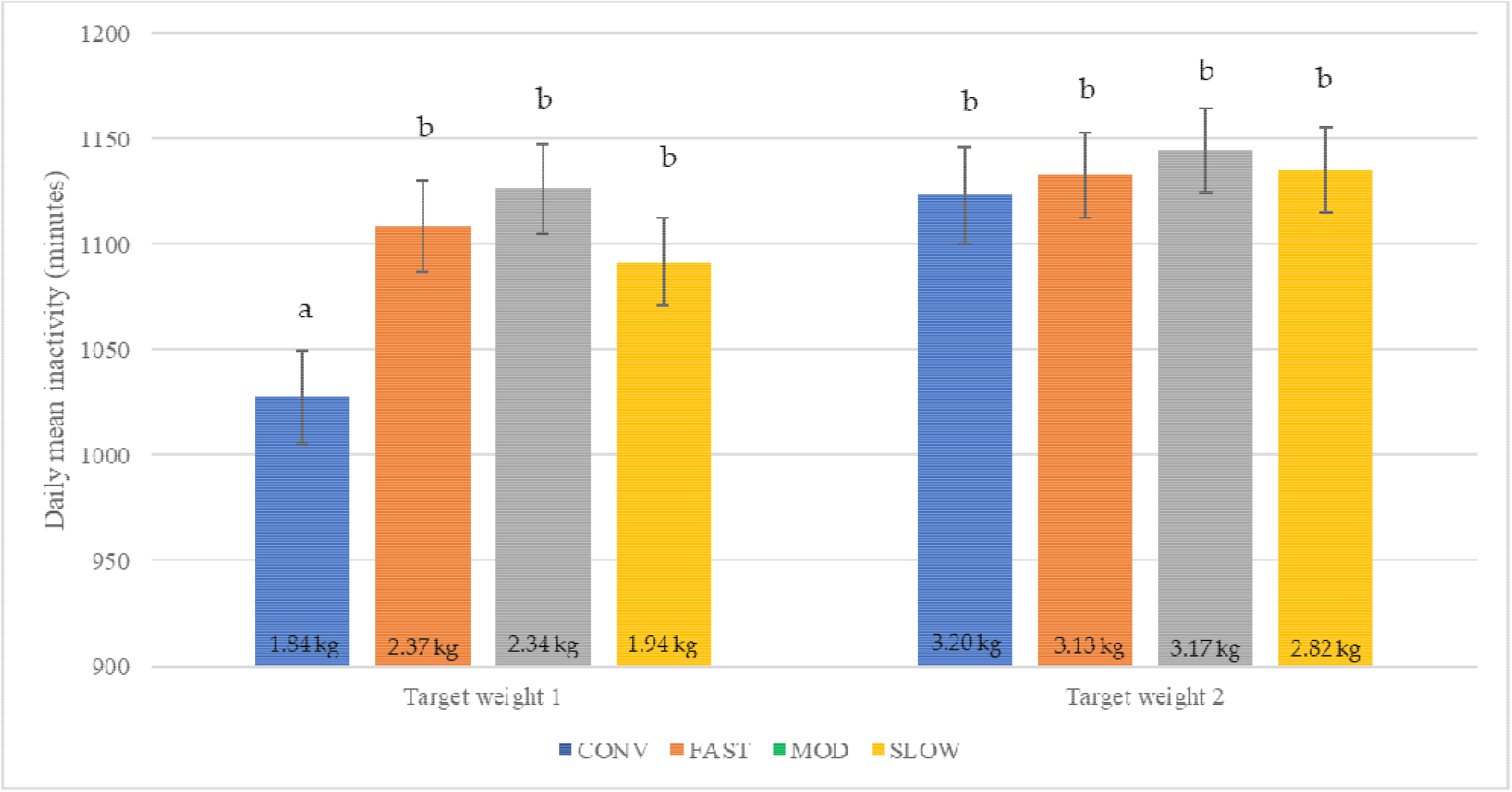
Mean daily inactivity levels (back-transformed LS-means ± SEM) by category at similar target weights. At target weight 1, CONV were 26 days of age, whereas the other strains were 42 days of age. At target weight 2, CONV were 42 days of age whereas the other strains were 56 days of age. Mean body weight (kg) within category is displayed within the bars. Bars with no common superscript differ at adjusted P < 0.05.

Inactivity levels were affected by the birds’ sex, with females again being more inactive than males (P < 0.0001). Females were inactive for 1127 ± 14 minutes per day, whereas males were inactive for 1094 ± 14 minutes per day, or 33 minutes less than females. The effect of sex on inactivity did not depend on target weight (F_1,187_ = 0.20, P = 0.6590), category (F_3, 187_ = 0.23, P = 0.8722), or strain nested within category (F_10, 187_ = 0.28, P = 0.9841).

### Behavioral Observations

A total of 1631 observations were analyzed, amounting to 678, 640, and 313 observations at days 26, 42, and 56, respectively. Analysis included data from 622 individual birds and 164 pens. Each strain was represented by 72–148 observations of 32–56 birds in 8–12 pens.

Some behaviours occurred too infrequently to analyze statistically. Foraging and dustbathing occurred in 4% and 2% of all observations, respectively, making the mean proportion of time spent foraging and dustbathing only 0.29% and 0.74% overall; when they did occur, they amounted to 8.26% and 35.5% of the observation. In terms of the event behaviours that occurred too infrequently to analyze, the following mean number of events were observed per 5-min observation period: wing flapping: 0.09; jumping: 0.04; feather ruffling: 0.10; head scratching: 0.42; pecking at rope: 0.11; and pecking at backpack: 0.01.

#### By Age

The time spent performing many behaviors was affected by category, but these differences only occurred at younger ages (Figure 3). At day 26, CONV spent more time sitting (Figure 3a) and less time standing (Figure 3b) and walking (Figure 3c) than FAST, MOD, and SLOW, but birds in all four categories spent similar amounts of time feeding (Figure 3d), drinking and preening. At day 42, CONV spent less time sitting than FAST, less time standing than MOD and SLOW, less time walking than SLOW, and more time feeding than FAST, MOD and SLOW. There were also differences between the slower-growing categories at day 42, with FAST sitting more and walking less than SLOW. Nonetheless, at day 42, there were no differences between the length of time any category spent drinking or preening. Additionally, at day 56, there were no differences between categories in the duration of any behaviors. At all ages, in terms of differences between categories, ‘all sitting’ was similar to sitting, ‘all standing’ was similar to standing, and locomotion was similar to walking; in other words, accounting for the time spent sidelying, performing other behaviors while sitting or standing, or running did not change the results. When behaviors were compiled to reflect the total time spent off feet, differences between categories only emerged at day 26, with CONV spending more time off feet than MOD and SLOW (Figure 3e).

**Figure 3.**
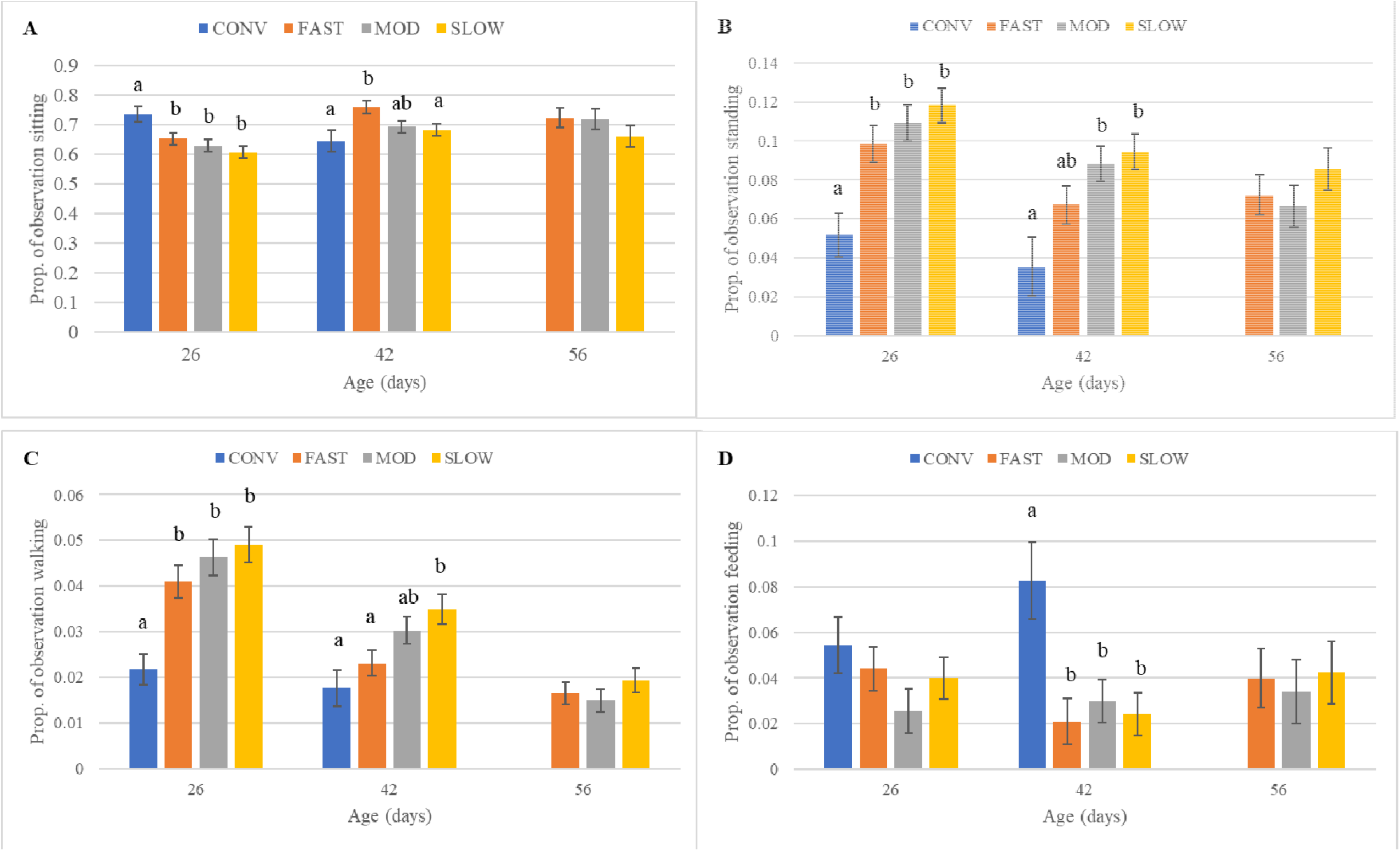
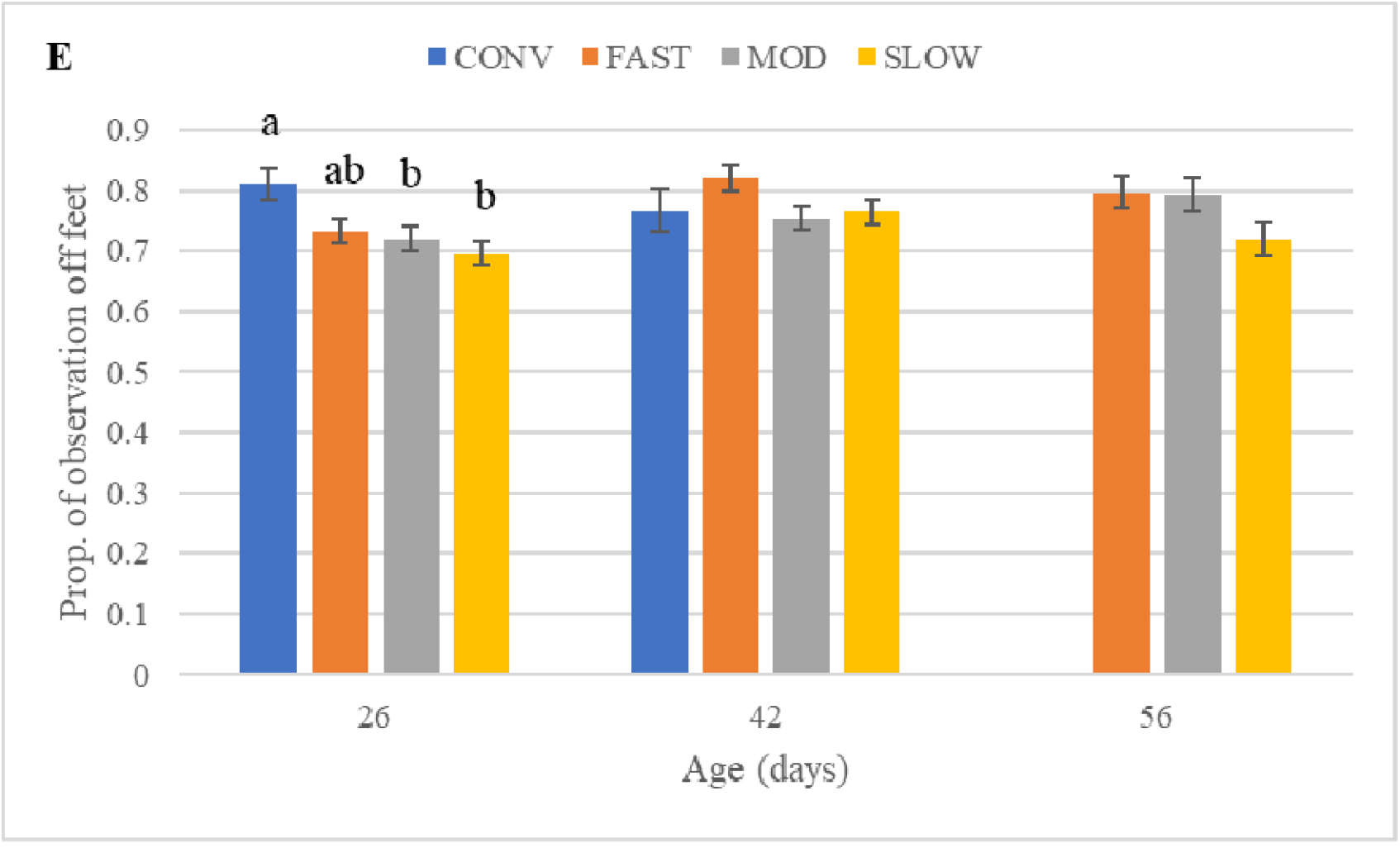
Proportion of time (back-transformed LS-means ± SEM) birds spent a) sitting; b) standing; c) walking; d) feeding; and e) off feet, assessed over a 300-second observation period, by category at the same age. Off feet is the sum duration of all sitting (including sitting while performing other behaviors, such as preening), sidelying, preening while sidelying, and dustbathing. Bars with no common superscript differ at adjusted P < 0.05.

Bout frequencies were also affected by category, although to a lesser extent than durations (Table 2). At day 26, CONV performed fewer bouts of standing and walking than all three slower-growing categories (i.e. FAST, MOD, and SLOW); in other words, it appears that CONV performed these behaviors less frequently overall, in addition to spending less time performing them (see above). Additionally, CONV engaged in fewer preening bouts than MOD and fewer leg stretches than FAST and MOD broilers. Similarly, at day 42, CONV also performed fewer bouts of standing and walking than MOD and SLOW, with FAST also performing fewer walking bouts than SLOW. There were no category differences in the number of bouts of any other behaviors at days 26 and 42, and no differences in any behaviors at day 56.

**Table 2.** Number of bouts (back-transformed LS-means ± SEM) of sitting, standing, walking, feeding, drinking, and preening, as well as number of events of pecking litter and leg stretching, as assessed over a 300-second live observation period, by category at the same age.

|  | CONV | FAST | MOD | SLOW |
| --- | --- | --- | --- | --- |
| **Day 26** |  |  |  |  |
| Sitting | 1.21 ± 0.08 | 1.14 ± 0.06 | 1.32 ± 0.06 | 1.21 ± 0.06 |
| Standing | 3.15 ± 0.71<sup>a</sup> | 5.70 ± 0.60<sup>b</sup> | 6.06 ± 0.59<sup>b</sup> | 5.76 ± 0.58<sup>b</sup> |
| Walking | 2.19 ± 0.29<sup>a</sup> | 3.72 ± 0.30<sup>b</sup> | 4.06 ± 0.32<sup>b</sup> | 3.63 ± 0.28<sup>b</sup> |
| Feeding | 0.68 ± 0.12 | 0.51 ± 0.10 | 0.38 ± 0.10 | 0.44 ± 0.09 |
| Drinking | 1.02 ± 0.50 | 1.71 ± 0.41 | 1.57 ± 0.41 | 1.54 ± 0.40 |
| Preening | 1.81 ± 0.27<sup>a</sup> | 2.10 ± 0.22<sup>ab</sup> | 2.68 ± 0.22<sup>b</sup> | 2.51 ± 0.21<sup>ab</sup> |
| Pecking litter | 3.27 ± 1.78 | 8.51 ± 1.45 | 6.02 ± 1.43 | 5.95 ± 1.38 |
| Leg stretching | 0.21 ± 0.06<sup>a</sup> | 0.49 ± 0.08<sup>b</sup> | 0.46 ± 0.08<sup>b</sup> | 0.39 ± 0.06<sup>ab</sup> |
| **Day 42** |  |  |  |  |
| Sitting | 0.91 ± 0.10 | 1.03 ± 0.07 | 1.02 ± 0.06 | 1.11 ± 0.06 |
| Standing | 2.13 ± 0.92<sup>a</sup> | 4.15 ± 0.63<sup>ab</sup> | 5.06 ± 0.59<sup>b</sup> | 5.20 ± 0.60<sup>b</sup> |
| Walking | 1.36 ± 0.30<sup>a</sup> | 2.13 ± 0.22<sup>ab</sup> | 2.78 ± 0.24<sup>ab</sup> | 3.11 ± 0.26<sup>b</sup> |
| Feeding | 0.61 ± 0.17 | 0.28 ± 0.10 | 0.33 ± 0.09 | 0.37 ± 0.10 |
| Drinking | 1.21 ± 0.65 | 1.53 ± 0.43 | 1.88 ± 0.40 | 1.64 ± 0.41 |
| Preening | 1.13 ± 0.35 | 1.72 ± 0.24 | 1.67 ± 0.22 | 1.75 ± 0.22 |
| Pecking litter | 3.34 ± 2.39 | 3.67 ± 1.52 | 6.16 ± 1.41 | 5.83 ± 1.43 |
| Leg stretching | 0.32 ± 0.11 | 0.19 ± 0.05 | 0.30 ± 0.06 | 0.30 ± 0.06 |
| **Day 56** |  |  |  |  |
| Sitting | - | 0.94 ± 0.08 | 0.99 ± 0.08 | 0.90 ± 0.08 |
| Standing | - | 3.22 ± 0.44 | 3.22 ± 0.45 | 3.83 ± 0.49 |
| Walking | - | 1.64 ± 0.23 | 1.63 ± 0.24 | 2.07 ± 0.28 |
| Feeding | - | 0.45 ± 0.15 | 0.44 ± 0.16 | 0.48 ± 0.16 |
| Drinking | - | 1.05 ± 0.34 | 1.12 ± 0.30 | 0.89 ± 0.28 |
| Preening | - | 1.19 ± 0.30 | 1.49 ± 0.32 | 1.95 ± 0.31 |
| Pecking litter | - | 6.25 ± 2.22 | 6.55 ± 2.36 | 7.32 ± 2.32 |
| Leg stretching | - | 0.22 ± 0.06 | 0.22 ± 0.06 | 0.28 ± 0.06 |

Within row, values with no common superscript differ at adj. $P < 0.05$

Within categories, strains generally behaved similarly, with a few exceptions. There were no differences between strains within CONV and MOD for both the total duration and number of bouts of all behaviors. Within FAST, there was one difference between strains: strain M stood for longer than strain F (13.76 ± 1.97% vs. 5.61 ± 1.49%, P = 0.0414), but only on day 26; these and all other FAST strains did not differ in their time budgets at any other ages (see Supplementary Table 4). Similarly, within SLOW, there were two differences between strains, and only at day 26: strain D spent a longer proportion of the observation walking than strain K (7.40 ± 0.90% vs. 3.43 ± 0.60%, P = 0.0231; see Supplementary Table 5) and strain J performed more sitting bouts than strain N (1.48 ± 0.11 vs. 1.02 ± 0.11, P = 0.0332); SLOW strains did not differ in their time budgets or bout frequencies at any other ages.

Behaviour was also affected by sex, but only for preening and feeding, with some interactions with category. In brief, at some ages and for some categories, females preened more and fed less than males (see Supplementary Material).

Finally, with respect to the time broilers spent on the litter, there were no differences between categories, with time ranging from 77.39% for MOD to 82.07% for CONV; however, there were differences between two strains within SLOW at day 26 (J = 87.65 ± 5.27% vs. K = 62.10 ± 5.46%, P = 0.0385) and between two strains within FAST at day 42 (F = 97.30 ± 5.04% vs. G = 71.60 ± 5.34%, P = 0.0215).

#### By Target Weight

At the same target weight, broilers in different categories behaved differently. At TW 1, CONV spent less time standing than MOD and SLOW, less time walking than SLOW, and more time feeding than FAST and SLOW (Figure 4a). FAST spent more time sitting than MOD and SLOW, and less time standing and walking than SLOW. Categories did not differ in the time spent drinking or preening. When compiled into off-feet behaviors, FAST spent more time off feet than MOD. By TW 2, almost all the category differences noted at TW 1 had disappeared. More specifically, categories did not differ in the length of time spent sitting, walking, feeding, drinking, or preening, nor did they differ in the total time spent off (Figure 4b); however, CONV did spend less time standing than MOD and SLOW (P = 0.0324 and 0.0028, respectively). As was noted for the age analysis, at both target weights, differences between categories in ‘all sitting’, ‘all standing’, and locomotion matched the differences noted in sitting, standing, and walking, respectively.

**Figure 4.**
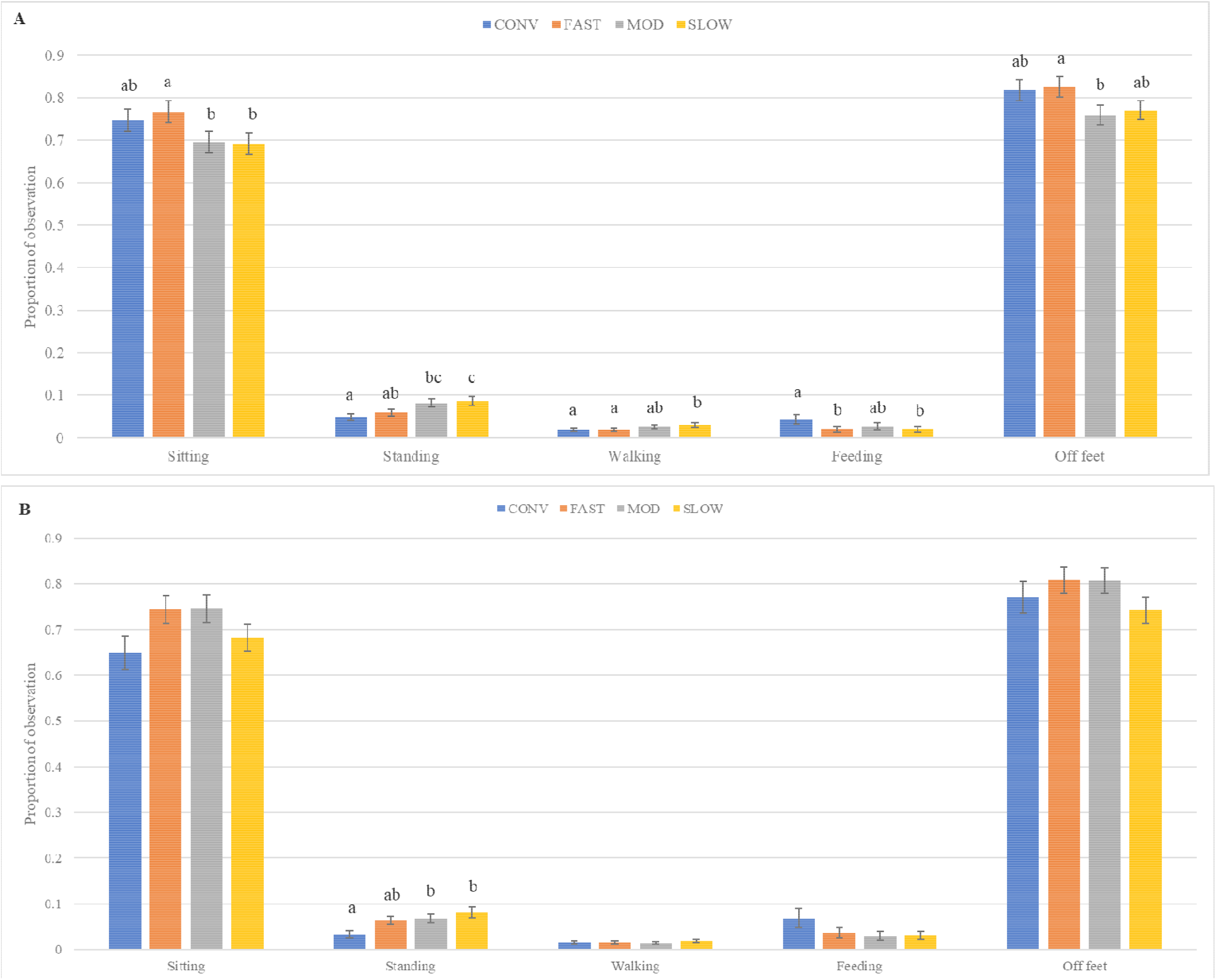
Proportion of time (back-transformed LS-means ± SEM) birds spent sitting, standing, walking, feeding, and off feet, assessed over a 300-second observation period, by category at a similar a) lighter target weight (target weight 1) and b) heavier target weight (target weight 2). Off feet is the sum duration of all sitting (including sitting while performing other behaviors, such as preening), sidelying, preening while sidelying, and dustbathing. At target weight 1, CONV were 26 days of age, whereas the other strains were 42 days of age. At target weight 2, CONV were 42 days of age whereas the other strains were 56 days of age. Bars with no common superscript differ at adjusted P < 0.05.

As shown in Table 3, bout frequencies were also affected by category, but only for some behaviors. At TW 1, there were differences across categories in the number of bouts of standing and walking, with CONV having fewer standing bouts than MOD and SLOW (both P < 0.05), and FAST having fewer walking bouts than SLOW (P = 0.0053). At TW 2, categories only differed in the number of preening bouts, with FAST performing fewer preening bouts than SLOW (P = 0.0463). Otherwise, at both target weights, there were no category differences in the bout frequencies of all other behaviors.

**Table 3.** Number of bouts (back-transformed LS-means ± SEM) of sitting, standing, walking, feeding, drinking, and preening, as well as number of events of pecking litter and leg stretching, as assessed over a 300-second live observation period, by category at similar target weights.

|  | CONV | FAST | MOD | SLOW |
| --- | --- | --- | --- | --- |
| **Target weight 1 (kg)** | 1.84 | 2.37 | 2.34 | 1.94 |
| Sitting | 1.17 ± 0.07 | 1.03 ± 0.06 | 1.01 ± 0.06 | 1.11 ± 0.06 |
| Standing | 3.10 ± 0.64<sup>a</sup> | 3.99 ± 0.62<sup>ab</sup> | 4.94 ± 0.59<sup>b</sup> | 5.07 ± 0.59<sup>b</sup> |
| Walking | 1.95 ± 0.32<sup>ab</sup> | 1.86 ± 0.30<sup>a</sup> | 2.42 ± 0.36<sup>ab</sup> | 2.72 ± 0.40<sup>b</sup> |
| Feeding | 0.68 ± 0.15 | 0.26 ± 0.14 | 0.29 ± 0.13 | 0.32 ± 0.09 |
| Drinking | 1.01 ± 0.54 | 1.62 ± 0.52 | 1.81 ± 0.50 | 1.76 ± 0.50 |
| Preening | 1.66 ± 0.25 | 1.53 ± 0.23 | 1.50 ± 0.21 | 1.51 ± 0.21 |
| Pecking litter | 3.05 ± 1.64 | 3.01 ± 1.54 | 6.33 ± 1.48 | 5.75 ± 1.47 |
| Leg stretching | 0.23 ± 0.06 | 0.20 ± 0.06 | 0.31 ± 0.06 | 0.29 ± 0.06 |
| **Target weight 2 (kg)** | 3.20 | 3.13 | 3.17 | 2.82 |
| Sitting | 0.97 ± 0.09 | 0.91 ± 0.07 | 0.98 ± 0.07 | 0.89 ± 0.07 |
| Standing | 2.22 ± 0.85 | 3.52 ± 0.72 | 3.50 ± 0.70 | 4.22 ± 0.71 |
| Walking | 1.23 ± 0.29 | 1.55 ± 0.29 | 1.52 ± 0.28 | 1.92 ± 0.33 |
| Feeding | 0.52 ± 0.19 | 0.50 ± 0.16 | 0.44 ± 0.16 | 0.50 ± 0.16 |
| Drinking | 1.31 ± 0.67 | 1.49 ± 0.59 | 1.52 ± 0.58 | 1.32 ± 0.58 |
| Preening | 1.04 ± 0.24<sup>ab</sup> | 1.01 ± 0.20<sup>a</sup> | 1.48 ± 0.24<sup>ab</sup> | 1.74 ± 0.28<sup>b</sup> |
| Pecking litter | 2.83 ± 2.27 | 6.20 ± 1.86 | 5.38 ± 1.84 | 6.81 ± 1.81 |
| Leg stretching | 0.29 ± 0.09 | 0.22 ± 0.07 | 0.20 ± 0.07 | 0.26 ± 0.07 |

Within row, values with no common superscript differ at adj. $P < 0.05$
At target weight 1, CONV were 26 days of age, whereas the other strains were 42 days of age.
At target weight 2, CONV were 42 days of age whereas the other strains were 56 days of age.
Mean body weight (kg) within category is displayed.

Similar to the pattern revealed through age analysis, at similar body weights, the strains within each category behaved similarly; there were no differences between strains within each of the four categories in terms of both the total duration and number of bouts for all analyzed behaviors (all P > 0.05).

Lastly, at the same target weight, there were no differences in the total duration each category spent on the litter, with values ranging from 80.71 ± 3.16% for CONV to 82.41 ± 2.97% for FAST.

### Enrichment Use

A total of 3167 scans, amounting to 1467, 1249, and 451 scans from days 26, 42, and 56, respectively, were performed. On each observation day, each pen was observed for 5–11 scans. Analysis included data from 163 pens. Each strain was represented in 8–12 pens.

#### By Age

The use of most enrichment objects was generally dependent on the category (Figure 5). At all ages, the proportion of birds using any of the available enrichments differed across categories (Figure 5a), with a lower proportion of CONV using all enrichments than all slower-growing categories. There were also differences amongst slower-growing categories: a higher proportion of SLOW than FAST and MOD birds used all enrichments on days 26 and 56. MOD enrichment use was also greater than FAST at day 56. A similar pattern was evident for the proportion of birds on the platform, with CONV always on the platform in a smaller proportion than all slower-growing categories, and some differences between slower-growing categories at day 26 and 56 (Figure 5b). Conversely, at all three ages, there were no differences in the proportion of birds using the area under the platform and ramp. Use of the hanging scale did not follow a consistent pattern in line with ADG (Figure 5c). MOD used the scale more than SLOW at day 26, and more than CONV at day 42. FAST used it less than MOD and SLOW at day 56. On the other hand, the proportion of birds on the litter generally decreased with decreasing ADG (Figure 5d), with all four categories using the litter differently on day 26, CONV differing from all slower-growing categories on day 42, and all three slower-growing categories differing from each other on day 56.

**Figure 5.**
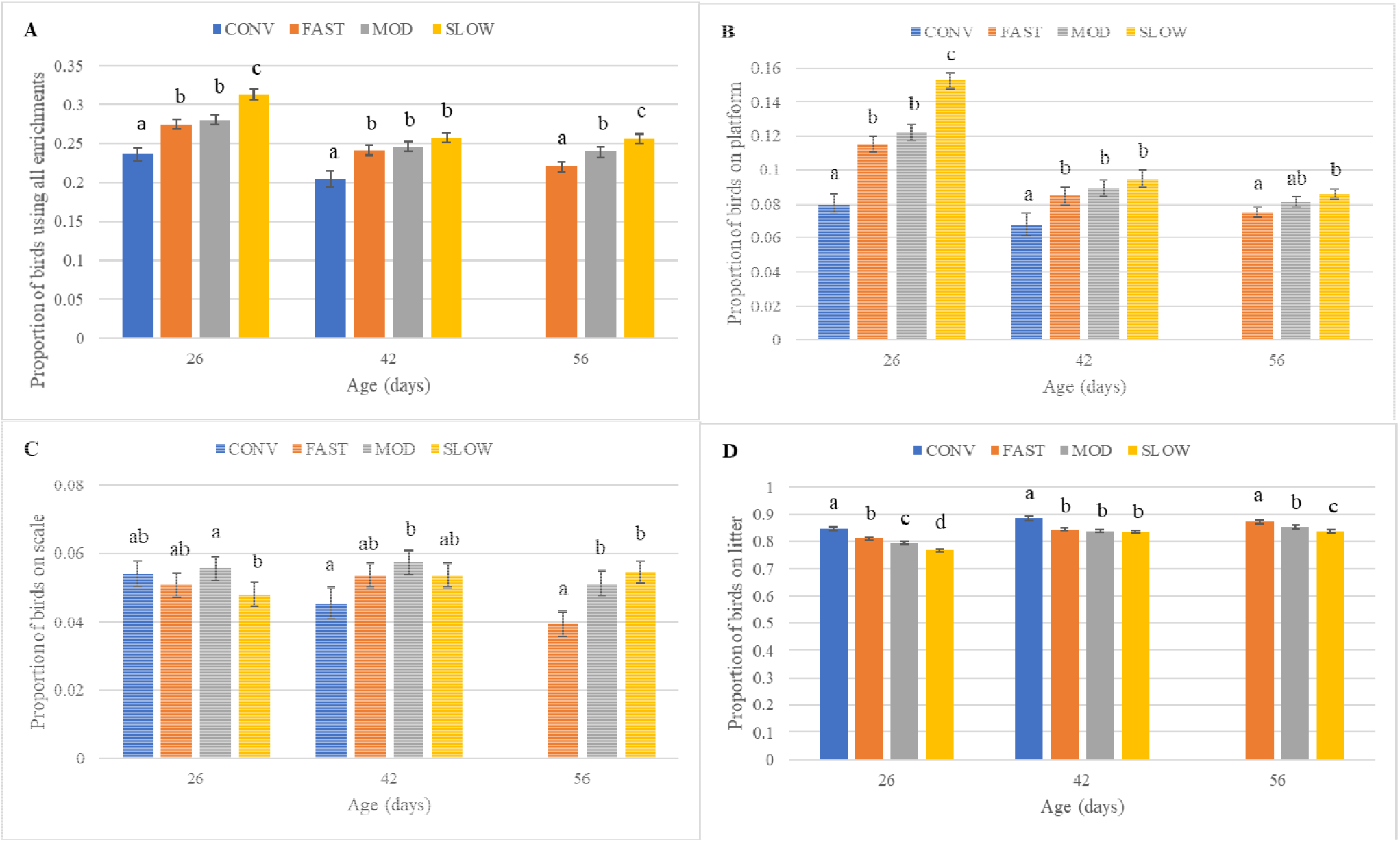
Enrichment use by category at the same age, assessed as the proportion of birds (back-transformed LS-means ± SEM) in the pen: a) using all enrichments; b) on the platform; c) on the scale; and d) on the litter. ‘All enrichments’ includes being on the platform, on the ramp, under the platform and ramp, on the scale, on and pecking the peck stone, and pecking the rope, whereas ‘on litter’ includes all birds not using any enrichment, but also includes those under the platform and ramp. Bars with no common superscript differ at adjusted P < 0.05.

Within categories, all CONV strains demonstrated a similar use of enrichments and the litter. Conversely, some strains within FAST, MOD, and SLOW used enrichments differently. Within the FAST and MOD categories (Supplementary Tables 6–7), enrichment use by one strain (F and E, respectively) was often different from some or all other strains within the category; however, within the SLOW category, there was no consistent pattern in the strains that differed from each other (Supplementary Table 8).

#### By Target Weight

At the same target body weight, enrichment use generally differed across categories, but not for every enrichment (Figure 6). At TW 1, a larger proportion of SLOW than CONV, FAST, and MOD were observed using all enrichments, on top of the platform, and using the area under the platform and ramp; however, there were no differences in the proportion of birds using the scale (Figure 6a). There were also few differences in the proportion of birds on the litter, with only FAST present on the litter in a greater proportion than SLOW (84.64 ± 0.56% vs. 83.46 ± 0.57%, P = 0.0012). At TW 2, the proportion of birds engaged in the use of enrichments generally increased as growth rate decreased (Figure 6b). CONV used all enrichments to a lesser extent than MOD and SLOW, whereas FAST and MOD each also used all enrichments less than SLOW. When each enrichment item was considered separately, there were fewer differences between categories: use of the top of the platform followed a similar pattern as all enrichment use, but with no difference between MOD and SLOW (P = 0.29), whereas there no differences in use of the area under the platform and ramp, and there were only differences in the use of the scale between the two faster (CONV, FAST) and two slower (MOD, SLOW) categories. Conversely, the proportion of birds on the litter decreased with growth rate, with a larger proportion of CONV on the litter (88.39 ± 0.78%) than all slower-growing categories (84.53 ± 0.49%, 83.84 ± 0.47%, and 83.59 ± 0.47% for FAST, MOD, and SLOW, respectively; all P < 0.03) and a greater proportion of FAST and MOD on the litter than SLOW (all P < 0.01).

**Figure 6.**
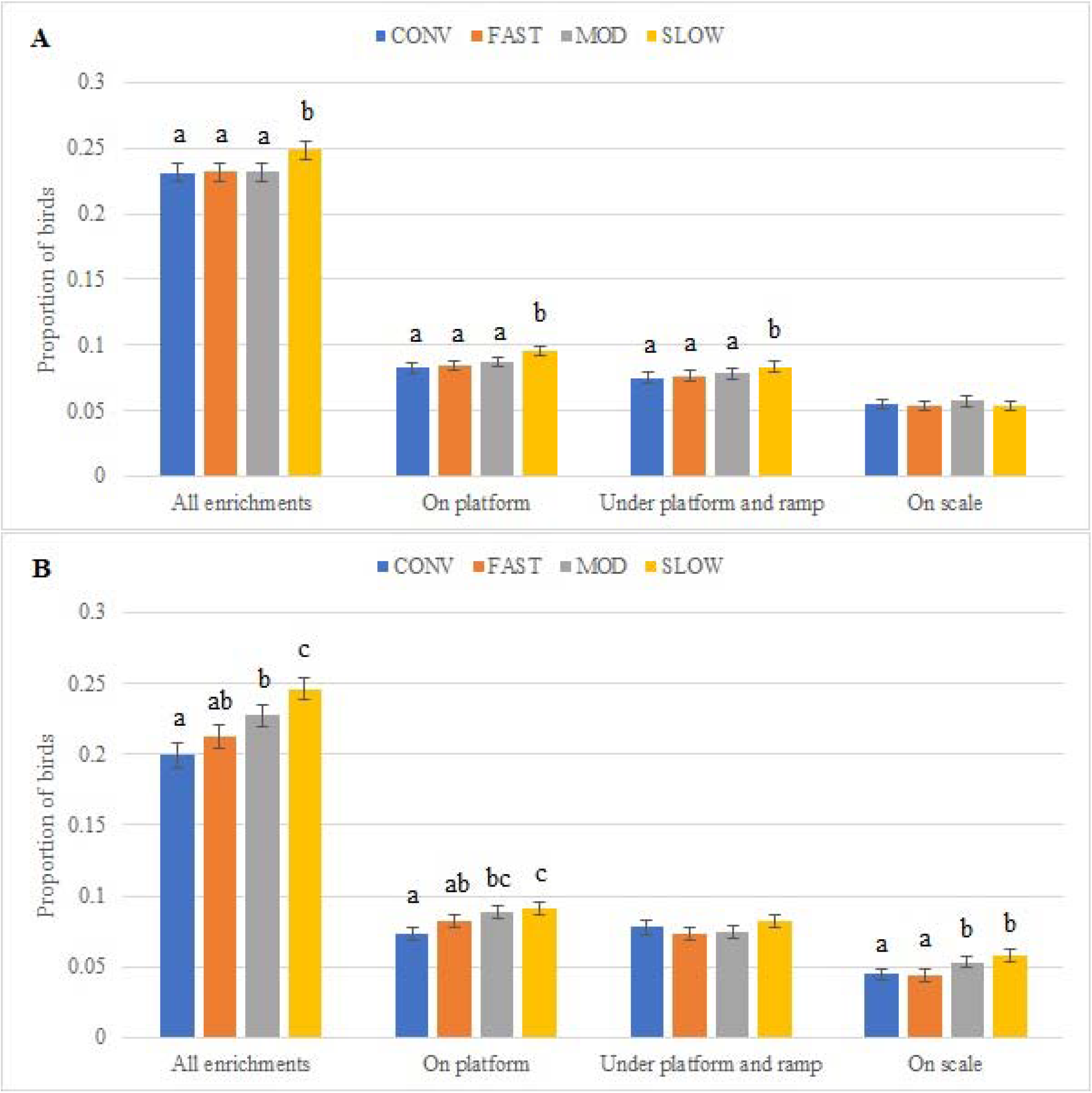
Enrichment use by category, assessed as the proportion of birds (back-transformed LS-means ± SEM) in the pen using various enrichment items and on the litter at a similar a) lighter target weight (target weight 1) and b) heavier target weight (target weight 2). ‘All enrichments’ includes being on the platform, on the ramp, under the platform and ramp, on the scale, on and pecking the peck stone, and pecking the rope, whereas ‘on litter’ includes all birds not using any enrichment, but also includes those under the platform and ramp. At target weight 1, CONV were 26 days of age, whereas the other strains were 42 days of age. At target weight 2, CONV were 42 days of age whereas the other strains were 56 days of age. Bars with no common superscript differ at adjusted P < 0.05.

Within categories, there were some differences in the number of birds that engaged in enrichment use. Amongst CONV, these differences only appeared at TW 1 and only in the proportion of birds on the platform (Supplementary Table 9). Within FAST, although each strain at some point demonstrated different enrichment use than the others, two strains were often different than the other FAST strains: F and I (Supplementary Table 10). Within MOD, strains E and O were the only strains to differ from each other, and only in their use of all enrichments and the scale (Supplementary Table 11). Lastly, within SLOW, enrichment use by each of the four strains was at some point different from the others, with no consistent pattern across enrichments (Supplementary Table 12).

## DISCUSSION

The aim of this study was to compare the behavior of conventional and slower-growing broiler strains, at both the same age and similar body weights, by investigating inactivity levels, behavior durations and frequencies, and enrichment use. All strains were inactive for most of the day, eventually reaching similar inactivity levels; however, those with slower growth rates reached these levels at older ages. At the same age, faster-growing strains had higher inactivity, but only at young ages. Behavioral time budgets were also affected by growth rate, with slower-growing strains sitting and feeding less, and standing and walking more than fast-growing strains; these differences were more widespread at younger ages and/or the lighter target weight. Enrichment use was higher in slower-growing strains. When compared at the same ages, the proportion of birds using enrichments was higher when their growth rates were lower, whereas the proportion of birds on the litter was higher when their growth rates were also higher. At both target weights, slower-growing strains engaged in more enrichment use; however, there were more differences between categories at the heavier body weight.

Surprisingly, at TW 1 (∼2.1 kg), CONV were less inactive than all three slower-growing categories, and spent a similar proportion of time sitting, as well as engaging in behaviors performed both off their feet. Additionally, at TW 1, CONV did not differ from FAST and MOD in enrichment use. CONV were two weeks younger than the other categories at TW 1 (34 vs. 48 days of age). However, CONV birds were also lighter than the expected target weight of 2.1 kg (∼1.8 kg at d34) and lighter than FAST and MOD (∼2.3 kg) birds at day 48 (although similar to SLOW birds, 1.9 kg, at d48; Torrey et al., 2021). This lighter weight may have been a consequence of consuming feed formulated for slower-growing strains or being processed 2 and 3 days prior to the breeders’ projections for reaching 2.1 kg; given the magnitude of the project, both adjustments were necessary for logistical reasons. At TW 2 (∼3.2 kg), CONV had a similar BW to FAST and MOD, but were heavier than SLOW (Torrey et al., 2021), suggesting that these mismatches in anticipated body weight was mainly an issue at the lower target weight. Nevertheless, given these weight differences, discussion of inactivity and behavior will focus on results from the age analyses. Conversely, discussion of enrichment use will include target weight comparisons due the intrinsic limitation of body size on the number of birds that could use each enrichment item.

As expected, all broiler strains eventually reached high levels of inactivity, equivalent to approximately 78–80% of the day, despite being provided with enrichment objects that may increase activity. Other research, based on behavioral observations, has shown that conventional birds spend 70–86% of their time performing inactive behaviors (e.g. Weeks et al., 2000; Bokkers and Koene, 2003; Alvino et al., 2009; Bailie et al., 2013; Dixon, 2020). Few studies have raised slower-growing broilers to 62 days of age; however, one study noted that older slower-growing broilers (> 7 weeks of age) spent 68% of observations sitting and lying (Bokkers and Koene, 2003). In contrast to previous research, our study continuously monitored activity, including overnight when broilers are known to be less active (Norring et al., 2016). As such, higher inactivity levels were expected compared to values from previous research; this expectation was also anticipated because sitting peaked at approximately 68–76% of observations, revealing that accelerometers capture more inactivity than day-time behavioral observations alone. On the other hand, the current results are likely conservative estimates of inactivity due to the accelerometer’s sensitivity to small movements and the methodology applied to define each inactivity count (i.e. 15 *consecutive* seconds of zero acceleration). This likely excluded bouts of low intensity, non-locomotor behaviors, such as preening, which has been shown to lead to non-zero accelerometer counts in laying hens (Casey-Trott and Widowski, 2018). It may have also excluded periods of rest during which a bird wearing an Actical^®^ pecked at the litter, was bumped or pecked at by pen mates, or shifted between resting postures (e.g. transitioning from sitting to sidelying).

Observational behavioral data identified differences in the performance of specific behaviors, thus providing information to supplement general inactivity results. For some behaviors, the time budgets noted in this study were similar to those noted in other studies of conventional and slower-growing broilers. For instance, across all strains and ages, the broilers in this study sat, stood, walked, and preened for about 60–75%, 3–12%, 1–5%, 5–10% of observations, whereas Bokkers and Koene (2003) and Dixon (2020) noted that sitting, standing, locomotion, and preening accounted for approximately 45–70%, 4–15%, 1–14%, and 4–12% of observations in similar-aged broilers. On the other hand, dustbathing and foraging were observed so infrequently that they could not be formally analyzed; they accounted for 0.74% and 0.29% of observations, compared to previous findings of 0–7% and 8–13%, respectively (Bokkers and Koene, 2003; Dixon, 2020). Dustbathing is known to be affected by time of day, occurring more frequently in the afternoon (De Jong and Gunnink, 2019); therefore, the comparatively low occurrence in this study may be due to the timing of our behavioral observations (morning to early afternoon). In terms of foraging, the definition used by Dixon (2020) included “scratching or digging…with the beak or feet”, whereas our definition required ground scratching with both legs in addition to ground pecking; our definition was thus more specific, and may have excluded instances that others may have considered to be foraging.

At the same age, platform use generally mimicked all enrichment use, suggesting that platforms were the most used enrichment; this may have been driven by a preference for elevated platforms over other types of enrichments, or by virtue of the platforms being large enough to simultaneously accommodate more birds than any other enrichment. At TW 2, when categories generally had similar body weights (Torrey et al., 2021), enrichment use followed growth rate, with faster-growing categories using enrichments less than slower-growing categories; this suggests that physical ability rather than body size itself limited platform use, particularly since access necessitated mobility: walking up an inclined ramp. Similarly, Rayner et al. (2020) noted that faster-growing broiler strains were observed on top of rectangular straw bales less frequently than slower-growing strains. Since domestic fowl generally prefer to rest and roost in elevated locations, enrichment use has been suggested to be an indicator of positive welfare (e.g. Rayner et al., 2020). Conversely, oral enrichment items (i.e. the rope and stone) were used infrequently, in line with previous research demonstrating that platforms are used more often than oral enrichments (e.g. Bach et al., 2019).

Generally, behavioral data corroborated inactivity data. More specifically, the inactivity levels of CONV and MOD birds increased until week 6, whereas those of FAST and SLOW birds increased until week 8. Similarly, as the birds aged, the proportion of time spent off feet (numerically) increased, whereas the proportion of time spent on feet decreased. These increases in inactivity and off-feet behaviors, like sitting, as well as decreases in on-feet behaviors, like standing and walking, may be due to an increased body weight or increasingly poor leg health, whereas category differences in the timing of these increases may have arisen due to their differences in growth rates. As broilers grow, their resting metabolic rate accounts for an increasing proportion of energy consumption, leaving less energy available for locomotor activity (Tickle et al., 2018), which likely, at least partially, contributed to increasing inactivity and off-feet behavior. Although this study did not assess either inactivity or behavior during the first three weeks of life, Bizeray (2000) noted that conventional and slower-growing strains did not display differences in lying, standing, or walking behavior until day 15, suggesting that it is unlikely that categories differed for long before the birds were outfitted with the accelerometers.

For the most part, at the same age, strains with similar growth rates behaved similarly. Within the CONV category, all strains demonstrated similar inactivity, behavioral time budgets, and enrichment use. Conversely, although only in a limited number of situations, some of the strains within all three slower-growing categories differed from each other. Within all three categories, strains with similar growth rates sometimes engaged in different enrichment use; however, there was no consistent pattern with respect to specific outlier strains or specific enrichment types. Within the FAST category, two strains differed from each other in their standing durations, but only at the youngest age, and not for any other behaviors. Within the MOD category, one strain (H) had lower inactivity levels, but only at a young age; investigation of the interaction between sex and strain nested within category revealed that this difference was likely driven by the male birds. Lastly, two SLOW strains differed in walking duration, whereas the other two SLOW strains performed a different number of sitting bouts; thus, all four demonstrated differences at some point. Mancinelli et al. (2020) noted that some organically-reared slower-growing strains demonstrated different degrees of static behavior, despite having similar growth rates. This suggests that aspects other than overall growth rate may impact behavior, such as temperament or strain-specific growth curves. For instance, strain H had a lower ADG than other MOD strains from approximately 17–35d (Torrey et al., 2021), which corresponded to the time period when strain H inactivity differed from that of other MOD strains. Future analyses using other data collected as part of this large-scale study will examine whether outlier strains differ in other characteristics that may explain the limited within-category behavior differences.

Although there were few differences between strains with similar growth rates (within categories), inactivity and behavior were not similar across all slower-growing strains. Among the 12 slower-growing strains included in this study, ADG to the higher target weight had a wide range, from 43.6 to 55.5 g/day (Torrey et al., 2021). At the same age, FAST birds (ADG_0-61_ = 53.5-55.5 g/day) were as inactive as CONV birds (ADG_0-47_= 66.0-68.7 g/day); this was the case at all ages. FAST birds were more inactive than MOD (ADG_0-61_ = 50.2-51.2 g/day) and SLOW (ADG_0-61_ = 43.6-47.7 g/day) birds, although only at young ages. At 42 days of age, FAST also stood and walked as little as CONV, both in duration and number of bouts, and spent similar amounts of time both off their feet, while also sitting more and walking less than SLOW.

Changes in behavior, particularly active behaviors such as standing or walking, may be the result of a lack of physical ability or a lack of motivation to be active or perform certain behaviors; distinguishing between these two potential causes is important for interpreting behavior changes with respect to welfare. Different strains may have different temperaments, which could affect their overall behavior. Since conventional strains have been selected for high feed efficiency, it is possible that they have been indirectly selected for low locomotor activity (Bizeray et al., 2000). As such, the behavior differences demonstrated in this study could be due to differences in the inherent characteristics of the tested strains. If this is the case, then there is less cause for concern with respect to welfare. On the other hand, Bokkers and Koene (2004) found that walking for a food reward is limited by motivation in slow-growing birds and physical ability in fast-growing birds. Their results suggest that immobility rather than motivation may be responsible for more inactivity and sitting, and less standing and walking among conventional broiler strains, or broilers with heavier body weights due to either their strain or age; this may impair broiler welfare through the inability to perform behaviors that they are motivated to perform.

This study demonstrated that inactivity, behavior, and enrichment use differ with broilers’ growth rates, with faster-growing strains demonstrating more inactivity, more sitting and feeding, less standing and walking, and lower rates of enrichment use. Future analyses will investigate if and how these behaviors correlate with other outcome measures of health and welfare, such as the performance of behaviors indicative of positive welfare (e.g. play behavior), mobility as assessed via latency-to-lie and obstacle tests, or health-related outcomes associated with different behaviors, particularly sitting and general inactivity, such as the incidence and severity of foot pad dermatitis or hock burn. Altogether, results suggest that not all slower-growing strains are equal, nor are they all different from conventional strains. As such, the use of any strain labelled as ‘slow growing’, without considering its specific growth rate, may not necessarily lead to improved welfare, at least not in terms of decreased inactivity, the performance of a full behavioral repertoire, and an increased use of enrichments.

## ACKNOWLEDGMENTS

We are grateful for the in-kind contributions from the Ontario Agri-Food Innovation Alliance, the anonymous breeding companies and Protekta, Inc. We are indebted to the Arkell Research Station staff for their assistance with ensuring the project’s success. We thank all our student assistants for their aid with accelerometer installation and behavioral observations (in alphabetical order): Alan Abdulkadar, Madeleine Browne, Veronica Cheng, Melanie Felker, Narissa Leslie, Nyasha Mombeshora, Siobhan Mellors, Quinn Rausch, Megan Weckworth, and Leah Wellard. This research was funded by Global Animal Partnership (grant number 053143) and the Canada First Research Excellence Fund (grant number 499028/499064).

