## Supplementary Material for "In pursuit of a better broiler: A comparison of the inactivity, behavior, and enrichment use of fast- and slower-growing broiler chickens"

**Supplementary Table 1.** Full ethogram for live behavioral observations.

| **Behavior** | **Description** |
| --- | --- |
| **State** |  |
| Sitting | Immobile, with entire breast touching the ground. Does not include sitting while engaged in other types of behavior (e.g. excludes sidelying, feeding while sitting). |
| Sidelying | Similar to sitting, but the chicken is lying on their side. One or both legs are usually stretched. |
| Standing | Immobile while supported on both legs, body not touching the ground. Does not include standing while engaged in other types of behavior (e.g. excludes feeding while standing, foraging). |
| Walking | Slow forward movement, using legs. Does not include other types of locomotion such as running, wing-assisted running or play fighting. |
| Running | Faster forward movement, using legs, but without wing involvement. |
| Wing-assisted running | Faster forward movement, using legs and wings. |
| Feed standing | Downward pecking in feeder while standing. |
| Feed sitting | Downward pecking in feeder while sitting. |
| Drinking | Pecking at nipple drinker. |
| Preen standing | Moving the beak through the feathers while standing. |
| Preen sidelying | Moving the beak through the feathers while sidelying. |
| Preen sitting | Moving the beak through the feathers while sitting. |
| Foraging | Ground scratching using both legs accompanied by pecking on the ground. |
| Play fight | Two birds, hopping and “chest bumping” (no touching required) while facing on another. Does not include forceful pecking. |
| Dust bathing | Vertical wing shakes in a lying position, often scooping litter onto the back of the body. The end point is when the bird stands up. |
| **Event** |  |
| Wing flapping | Flapping wings while standing. |
| Jumping | Push off a surface and into the air using leg muscles. |
| Peck at stone | Peck at any part of the peck stone. |
| Peck at rope | Peck at any part of the rope. |
| Peck at backpack | Peck at any part of the Actical or straps. |
| Peck at litter | Peck at litter, no ground scratching involved. |
| Feather ruffle | Feather erection and body shaking. |
| Leg stretching | Stretching one of the legs while standing (sometimes accompanied by wing stretching). |
| Head scratching | Scratching head using one foot. |
| **Location** |  |
| On platform | On the horizontal part of the platform. |
| On ramp | On the graded slope of the ramp. |
| Under platform/ramp | Under the platform or ramp. |
| On scale | On the hanging scale. |
| On stone | On the pecking stone. |
| On drinker line | Perching on the drinker line. |
| On litter | On the litter, in any location other than those listed above. |

**RESULTS**

***Inactivity***

***By Age.*** Females were generally more inactive than males, but sex differences depended on the category. As shown in Supplementary Table 2, there was an interaction between sex and category, but only in the analysis for weeks 4-7 (F_3,6877_ = 4.28, P = 0.0050). Among female birds, inactivity levels were similar across all categories (all P > 0.05). Among male birds, CONV (1069 ± 23 min) were more inactive than SLOW (995 ± 13 min, P = 0.0253), but inactivity was similar across all other categories (all P > 0.05). Within each category, females were consistently more inactive than males (all P < 0.02).

There was also an interaction between sex and strain nested within category (Supplementary Table 3). At 4-7 weeks of age, the males and females from 7 out of 14 strains (1 CONV strain: C; 2 FAST strains: G, I; 2 MOD strains: H, O; 2 SLOW strains: J, N) had different inactivity levels (all P < 0.02); in these instances, females were more inactive than males. Females across strains within categories showed similar inactivity levels, as did males across CONV, FAST, and SLOW strains; however, H males had lower inactivity levels than the other MOD strain males (E, O, S). At 8-9 weeks of age, there was an effect of sex for 10 out of 12 strains (3 FAST strains: F, G, I; all 4 MOD strains: E, H, O, S; 3 SLOW strains: D, J, K); again, for these strains, females were more inactive than males. At 8-9 weeks of age, females across strains within categories showed similar inactivity levels, as did males (all P > 0.05).

***Behavioral Observations***

***By Age.*** Behavior was also affected by sex, but only for preening and feeding. At both days 26 and 42, females spent more time preening than males (10.65 ± 1.04 vs. 5.06 ± 1.03%, P < 0.0001), and performed more preening bouts (2.67 ± 0.22 vs. 1.59 ± 0.21, P < 0.001), but only amongst SLOW birds. At day 56, all females, regardless of category, spent more time preening than males (9.08 ± 1.08 vs. 3.56 ± 1.05%, P < 0.0001), as well as performed more preening bouts (2.02 ± 0.26 vs. 1.07 ± 0.26, P = 0.0008). Moreover, at day 56, females also spent less time feeding than males (2.35 ± 1.12 vs. 5.41 ± 1.08%, P = 0.0205).

**Supplementary Table 2.** The effect of category and sex on mean daily inactivity levels (back-transformed LS-means ± SEM) of broiler chickens at the same age, as measured using an omni-directional accelerometer (Actical^®^).

| **Category** | **Mean daily inactivity (min)** | |
| --- | --- | --- |
|  | Female | Male |
| **Weeks 4-7 (day 22-49)** | | |
| CONV | 1117.07 ± 24.09^a^ | 1068.96±23.25^by^ |
| FAST | 1074.75 ± 14.64^a^ | 1045.46 ± 14.72^b^ |
| MOD | 1080.91 ± 13.86^a^ | 1007.32 ± 13.01^b^ |
| SLOW | 1046.74 ± 14.11^a^ | 995.01 ± 13.20^bz^ |
| **Weeks 8-9 (day 50-60)** | | |
| FAST | 1163.77 ± 16.74^a^ | 1127.38 ± 16.40^b^ |
| MOD | 1164.38 ± 15.97^a^ | 1124.48 ± 15.45^b^ |
| SLOW | 1157.87 ± 16.87^a^ | 1129.62 ± 16.42^b^ |

^a ≠ b^ within rows at adjusted P < 0.05

^y ≠ z^ within columns at the same age at adjusted P < 0.05

**Supplementary Table 3.** The effect of strain nested within category and sex on mean daily inactivity levels (back-transformed LS-means ± SEM) of broiler chickens at the same age, as measured using an omni-directional accelerometer (Actical^®^).

| **Category** | **Strain** | **Mean daily inactivity (min)** | |
| --- | --- | --- | --- |
|  |  | Female | Male |
| **Weeks 4-7 (day 22-49)** | | | |
| CONV | B | 1081.62 ± 29.36 | 1056.48 ± 29.28 |
|  | C | 1153.67 ± 38.67^a^ | 1081.58 ± 36.26^b^ |
| FAST | F | 1078.33 ± 28.66 | 1070.73 ± 28.44 |
|  | G | 1099.88 ± 27.57^a^ | 1060.77 ± 26.57^b^ |
|  | I | 1093.28 ± 28.61^a^ | 1051.87 ± 29.40^b^ |
|  | M | 1028.96 ± 31.72 | 1000.62 ± 32.52 |
| MOD | E | 1099.71 ± 27.96 | 1069.39 ± 27.66^y^ |
|  | H | 1027.84 ± 26.97^a^ | 848.62 ± 22.60^bz^ |
|  | O | 1099.71 ± 28.41^a^ | 1049.66 ± 26.65^by^ |
|  | S | 1098.40 ± 27.50 | 1080.86 ± 27.02^y^ |
| SLOW | D | 1011.40 ± 28.32 | 982.16 ± 27.02 |
|  | J | 1073.18 ± 28.54^a^ | 1030.41 ± 27.40^b^ |
|  | K | 1028.63 ± 26.12 | 1001.25 ± 25.42 |
|  | N | 1075.24 ± 29.83^a^ | 967.34 ± 25.76^b^ |
| **Weeks 8-9 (day 50-60)** | | | |
| FAST | F | 1199.31 ± 31.58^a^ | 1138.76 ± 30.12^b^ |
|  | G | 1185.71 ± 32.45^a^ | 1166.87 ± 31.93^b^ |
|  | I | 1170.68 ± 32.61^a^ | 1111.59 ± 31.12^b^ |
|  | M | 1101.68 ± 37.29 | 1093.66 ± 38.02 |
| MOD | E | 1175.15 ± 36.57^a^ | 1152.69 ± 35.30^b^ |
|  | H | 1115.96 ± 28.57^a^ | 1054.45 ± 26.97^b^ |
|  | O | 1165.38 ± 32.01^a^ | 1121.61 ± 30.64^b^ |
|  | S | 1202.75 ± 32.85^a^ | 1172.82 ± 32.03^b^ |
| SLOW | D | 1147.23 ± 31.25^a^ | 1097.63 ± 29.74^b^ |
|  | J | 1139.44 ± 35.01^a^ | 1107.32 ± 34.02^b^ |
|  | K | 1178.77 ± 32.38 | 1170.38 ± 31.97 |
|  | N | 1166.44 ± 35.74^a^ | 1144.65 ± 35.07^b^ |

^a ≠ b^ within rows at adjusted P < 0.05

^y ≠ z^ within columns at the same age at adjusted P < 0.05

**Supplementary Table 4.** Proportion of time individual birds spent performing various behaviors (back-transformed LS-means ± SEM), assessed over a 300-second live observation period, within FAST strains according to bird age.

|  | **F** | **G** | **I** | **M** |
| --- | --- | --- | --- | --- |
| **Day 26** |  |  |  |  |
| Sitting (%) | 67.63 ± 3.53 | 68.95 ± 3.80 | 63.05 ± 3.83 | 61.59 ± 4.69 |
| Standing (%) | 5.61 ± 1.49^a^ | 9.61 ± 1.60^ab^ | 10.38 ± 1.61^ab^ | 13.76 ± 1.97^b^ |
| Walking (%) | 3.88 ± 0.62 | 3.33 ± 0.59 | 4.07 ± 0.64 | 5.35 ± 0.98 |
| Feeding (%) | 4.90 ± 1.64 | 4.43 ± 1.79 | 6.22 ± 1.79 | 2.13 ± 2.19 |
| Drinking (%) | 4.17 ± 1.25 | 2.98 ± 1.19 | 5.97 ± 1.74 | 7.60 ± 2.44 |
| Preening (%) | 6.98 ± 1.65 | 6.45 ± 1.77 | 6.87 ± 1.78 | 9.83 ± 2.18 |
| Off feet (%) | 76.72 ± 3.46 | 77.60 ± 3.75 | 70.71 ± 3.76 | 68.09 ± 4.60 |
| On feet (%) | 19.32 ± 3.12 | 22.14 ± 3.50 | 28.69 ± 3.89 | 32.50 ± 5.07 |
| **Day 42** |  |  |  |  |
| Sitting (%) | 80.69 ± 3.64 | 74.94 ± 3.81 | 75.43 ± 3.88 | 72.68 ± 4.82 |
| Standing (%) | 4.12 ± 1.54 | 6.15 ± 1.60 | 6.91 ± 1.63 | 9.69 ± 2.01 |
| Walking (%) | 2.21 ± 0.47 | 2.02 ± 0.48 | 2.42 ± 0.51 | 2.66 ± 0.71 |
| Feeding (%) | 1.51 ± 1.68 | 0.80 ± 1.79 | 1.68 ± 1.80 | 4.45 ± 2.26 |
| Drinking (%) | 3.80 ± 1.23 | 5.77 ± 1.70 | 6.86 ± 1.91 | 4.29 ± 1.77 |
| Preening (%) | 3.79 ± 1.71 | 6.74 ± 1.77 | 5.60 ± 1.81 | 6.88 ± 2.22 |
| Off feet (%) | 87.65 ± 3.57 | 83.77 ± 3.75 | 80.40 ± 3.81 | 75.99 ± 4.73 |
| On feet (%) | 12.56 ± 2.58 | 16.16 ± 3.08 | 19.42 ± 3.38 | 24.06 ± 4.75 |
| **Day 56** |  |  |  |  |
| Sitting (%) | 77.72 ± 5.30 | 70.04 ± 6.12 | 73.48 ± 6.46 | 68.16 ± 8.50 |
| Standing (%) | 4.58 ± 1.67 | 7.25 ± 2.06 | 5.08 ± 2.12 | 11.99 ± 2.74 |
| Walking (%) | 1.49 ± 0.36 | 1.15 ± 0.40 | 1.48 ± 0.44 | 2.91 ± 0.93 |
| Feeding (%) | 2.75 ± 2.23 | 4.11 ± 2.59 | 2.52 ± 2.72 | 6.57 ± 3.55 |
| Drinking (%) | 8.19 ± 2.68 | 8.36 ± 3.31 | 8.61 ± 3.40 | 7.38 ± 4.40 |
| Preening (%) | 2.67 ± 1.99 | 5.79 ± 2.31 | 4.79 ± 2.38 | 5.89 ± 3.16 |
| Off feet (%) | 80.70 ± 4.40 | 75.98 ± 5.42 | 83.33 ± 5.58 | 78.95 ± 7.21 |
| On feet (%) | 17.59 ± 5.57 | 22.15 ± 7.46 | 15.42 ± 6.32 | 24.38 ± 10.66 |

^a ≠ b^ within rows at adjusted P < 0.05

**Supplementary Table 5.** Proportion of time individual birds spent performing various behaviors (back-transformed LS-means ± SEM), assessed over a 300-second live observation period, within SLOW strains according to bird age.

|  | **D** | **J** | **K** | **N** | **T^1^** |
| --- | --- | --- | --- | --- | --- |
| **Day 26** |  |  |  |  |  |
| Sitting (%) | 53.07 ± 3.58 | 62.50 ± 3.76 | 64.85 ± 3.87 | 62.85 ± 3.82 | 27.12 ± 8.12 |
| Standing (%) | 14.95 ± 1.53 | 9.97 ± 1.58 | 11.81 ± 1.62 | 10.59 ± 1.60 | 33.33 ± 6.76 |
| Walking (%) | 7.40 ± 0.90^a^ | 4.46 ± 0.67^ab^ | 3.43 ± 0.60^b^ | 5.07 ± 0.75^ab^ | 14.05 ± 3.29 |
| Feeding (%) | 5.88 ± 1.67 | 6.74 ± 1.77 | 0.00 ± 1.82 | 4.22 ± 1.80 | 3.14 ± 2.53 |
| Drinking (%) | 4.58 ± 1.45 | 5.80 ± 1.63 | 2.85 ± 1.23 | 6.83 ± 1.80 | 1.87 ± 0.92 |
| Preening (%) | 7.97 ± 1.72 | 9.18 ± 1.75 | 12.10 ± 1.80 | 5.66 ± 1.77 | 8.00 ± 1.81 |
| Off feet (%) | 61.07 ± 3.50 | 70.85 ± 3.69 | 76.63 ± 3.81 | 69.80 ± 3.75 | 40.85 ± 9.05 |
| On feet (%) | 40.32 ± 4.11 | 28.46 ± 3.82 | 23.04 ± 3.60 | 29.42 ± 3.95 | 59.15 ± 9.05 |
| **Day 42** |  |  |  |  |  |
| Sitting (%) | 60.59 ± 3.58 | 73.57 ± 3.90 | 68.99 ± 3.86 | 70.13 ± 3.80 | 30.97 ± 6.65 |
| Standing (%) | 10.31 ± 1.54 | 8.53 ± 1.64 | 10.09 ± 1.62 | 8.81 ± 1.60 | 32.83 ± 4.76 |
| Walking (%) | 4.05 ± 0.64 | 2.97 ± 0.57 | 3.23 ± 0.59 | 3.85 ± 0.67 | 13.98 ± 1.59 |
| Feeding (%) | 4.41 ± 1.67 | 1.53 ± 1.83 | 1.94 ± 1.82 | 1.88 ± 1.78 | 4.18 ± 2.47 |
| Drinking (%) | 2.89 ± 1.06 | 5.20 ± 1.63 | 6.60 ± 1.92 | 6.19 ± 1.71 | 2.97 ± 1.05 |
| Preening (%) | 7.39 ± 1.72 | 5.13 ± 1.81 | 7.04 ± 1.80 | 8.36 ± 1.77 | 11.68 ± 4.30 |
| Off feet (%) | 73.04 ± 3.51 | 79.81 ± 3.83 | 76.27 ± 3.80 | 76.74 ± 3.73 | 34.46 ± 6.93 |
| On feet (%) | 26.88 ± 3.66 | 20.01 ± 3.46 | 23.34 ± 3.63 | 22.88 ± 3.61 | 65.54 ± 6.93 |
| **Day 56** |  |  |  |  |  |
| Sitting (%) | 60.01 ± 5.76 | 70.76 ± 6.13 | 63.85 ± 6.67 | 70.15 ± 6.20 | 32.77 ± 6.86 |
| Standing (%) | 11.64 ± 1.77 | 6.43 ± 2.01 | 7.17 ± 2.50 | 9.02 ± 2.04 | 32.70 ± 5.71 |
| Walking (%) | 3.04 ± 0.60 | 1.59 ± 0.41 | 1.71 ± 0.48 | 1.73 ± 0.46 | 10.10 ± 1.87 |
| Feeding (%) | 1.12 ± 2.32 | 6.97 ± 2.56 | 6.32 ± 2.87 | 2.56 ± 2.60 | 1.71 ± 1.45 |
| Drinking (%) | 11.27 ± 2.84 | 6.06 ± 3.22 | 7.51 ± 3.61 | 10.24 ± 3.28 | 4.23 ± 1.36 |
| Preening (%) | 10.71 ± 2.18 | 8.29 ± 2.26 | 6.86 ± 2.52 | 5.41 ± 2.30 | 15.11 ± 4.00 |
| Off feet (%) | 64.42 ± 4.65 | 77.45 ± 5.29 | 75.36 ± 5.92 | 70.76 ± 5.38 | 35.79 ± 7.27 |
| On feet (%) | 34.82 ± 7.80 | 21.18 ± 6.95 | 23.01 ± 8.08 | 27.65 ± 7.97 | 64.21 ± 7.27 |

^1^ Raw means are presented; due to a low sample size, strain T (ADG_0-61_ = 19.78 g/day) was not included in the statistical models.

^a ≠ b^ within rows at adjusted P < 0.05

**Supplementary Table 6.** Enrichment use, assessed as the percentage of birds in the pen using each enrichment (back-transformed LS-means ± SEM), within FAST strains according to bird age.

|  | **F** | **G** | **I** | **M** |
| --- | --- | --- | --- | --- |
| **Day 26** |  |  |  |  |
| All enrichments (%) | 25.25 ± 1.14 | 30.06 ± 1.05 | 30.08 ± 1.15 | 24.25 ± 1.49 |
| On platform (%) | 8.42 ± 0.74^a^ | 13.63 ± 0.71^b^ | 13.39 ± 0.74^b^ | 10.66 ± 0.93^ab^ |
| Under platform and ramp (%) | 11.46 ± 0.96^a^ | 8.86 ± 0.90^ab^ | 8.01 ± 0.96^ab^ | 5.94 ± 1.21^b^ |
| On scale (%) | 3.85 ± 0.47^a^ | 5.17 ± 0.46^ab^ | 5.88 ± 0.47^b^ | 5.41 ± 0.57^ab^ |
| On litter (%) | 85.75 ± 0.82^a^ | 78.91 ± 0.75^b^ | 78.05 ± 0.08^b^ | 81.13 ± 1.03^b^ |
| **Day 42** |  |  |  |  |
| All enrichments (%) | 20.41 ± 1.18^a^ | 26.03 ± 1.09^b^ | 27.09 ± 1.15^b^ | 22.98 ± 1.39^ab^ |
| On platform (%) | 4.90 ± 0.77^a^ | 9.42 ± 0.74^b^ | 9.87 ± 0.76^b^ | 9.78 ± 0.89^b^ |
| Under platform and ramp (%) | 10.42 ± 0.99 | 9.35 ± 0.93 | 9.17 ± 0.98 | 5.86 ± 1.17 |
| On scale (%) | 4.56 ± 0.49 | 5.66 ± 0.47 | 6.19 ± 0.48 | 4.98 ± 0.54 |
| On litter (%) | 89.66 ± 0.84^a^ | 83.53 ± 0.79^b^ | 82.23 ± 0.80^b^ | 82.71 ± 0.96^b^ |
| **Day 56** |  |  |  |  |
| All enrichments (%) | 19.03 ± 0.99^a^ | 24.53 ± 0.93^b^ | 26.13 ± 0.94^b^ | 18.33 ± 1.43^a^ |
| On platform (%) | 5.45 ± 0.48^a^ | 9.14 ± 0.45^b^ | 8.03 ± 0.45^b^ | 7.39 ± 0.70^ab^ |
| Under platform and ramp (%) | 9.26 ± 0.62^a^ | 10.50 ± 0.58^a^ | 10.42 ± 0.58^a^ | 5.73 ± 0.91^b^ |
| On scale (%) | 3.40 ± 0.54 | 3.67 ± 0.50 | 5.61 ± 0.51 | 3.03 ± 0.78 |
| On litter (%) | 90.27 ± 0.85^a^ | 85.85 ± 0.81^b^ | 84.23 ± 0.83^b^ | 88.12 ± 1.31^ab^ |

^a ≠ b^ within rows at adjusted P < 0.05

**Supplementary Table 7.** Enrichment use, assessed as the percentage of birds in the pen using each enrichment (back-transformed LS-means ± SEM), within MOD strains according to bird age.

|  | **E** | **H** | **O** | **S** |
| --- | --- | --- | --- | --- |
| **Day 26** |  |  |  |  |
| All enrichments (%) | 24.12 ± 1.17^a^ | 28.06 ± 1.20^ab^ | 29.72 ± 10.8^b^ | 30.39 ± 1.16^b^ |
| On platform (%) | 11.19 ± 0.76^ab^ | 12.44 ± 0.80^ab^ | 11.21 ± 0.73^a^ | 13.96 ± 0.77^b^ |
| Under platform and ramp (%) | 7.08 ± 0.99 | 7.68 ± 1.02 | 9.35 ± 0.93 | 7.06 ± 0.99 |
| On scale (%) | 3.92 ± 0.47^a^ | 5.30 ± 0.50^ab^ | 6.30 ± 0.47^b^ | 6.72 ± 0.48^b^ |
| On litter (%) | 82.59 ± 0.81^a^ | 78.79 ± 0.86^b^ | 79.80 ± 0.77^a^ | 76.84 ± 0.80^b^ |
| **Day 42** |  |  |  |  |
| All enrichments (%) | 21.05 ± 1.20^a^ | 24.66 ± 1.17^ab^ | 26.97 ± 1.10^b^ | 25.66 ± 1.17^ab^ |
| On platform (%) | 7.46 ± 0.78 | 9.10 ± 0.77 | 9.48 ± 0.75 | 9.70 ± 0.78 |
| Under platform and ramp (%) | 7.64 ± 1.01 | 8.49 ± 1.00 | 9.13 ± 0.94 | 8.52 ± 1.01 |
| On scale (%) | 4.81 ± 0.49 | 5.43 ± 0.48 | 6.53 ± 0.48 | 6.15 ± 0.49 |
| On litter (%) | 86.36 ± 0.84^a^ | 83.60 ± 0.81^ab^ | 82.33 ± 0.79^b^ | 83.08 ± 0.82^ab^ |
| **Day 56** |  |  |  |  |
| All enrichments (%) | 21.75 ± 1.03^a^ | 21.79 ± 1.22^a^ | 27.24 ± 096^b^ | 24.89 ± 0.98^ab^ |
| On platform (%) | 7.18 ± 0.51 | 7.37 ± 0.58 | 9.29 ± 0.47 | 8.70 ± 0.47 |
| Under platform and ramp (%) | 8.58 ± 0.65 | 7.95 ± 0.76 | 10.30 ± 0.60 | 10.56 ± 0.61 |
| On scale (%) | 4.24 ± 0.56 | 5.58 ± 0.66 | 5.93 ± 0.52 | 4.71 ± 0.53 |
| On litter (%) | 86.71 ± 0.95 | 86.22 ± 1.12 | 83.19 ± 0.84 | 85.81 ± 0.86 |

^a ≠ b^ within rows at adjusted P < 0.05

**Supplementary Table 8.** Enrichment use, assessed as the percentage of birds in the pen using each enrichment (back-transformed LS-means ± SEM), within SLOW strains according to bird age.

|  | **D** | **J** | **K** | **N** | **T^1^** |
| --- | --- | --- | --- | --- | --- |
| **Day 26** |  |  |  |  |  |
| All enrichments (%) | 27.78 ± 1.19^a^ | 32.63 ± 1.08^ab^ | 34.35 ± 1.12^b^ | 30.54 ± 1.10^ab^ | 30.72 ± 0.45 |
| On platform (%) | 13.46 ± 0.79 | 15.22 ± 0.71 | 16.25 ± 0.76 | 16.11 ± 0.73 | 15.99 ± 2.33 |
| Under platform and ramp (%) | 5.90 ± 1.01^a^ | 10.93 ± 0.93^b^ | 8.68 ± 0.98^ab^ | 7.75 ± 0.95^ab^ | 2.76 ± 0.40 |
| On scale (%) | 4.98 ± 0.49^ab^ | 4.33 ± 0.45^a^ | 6.13 ± 0.78^b^ | 3.77 ± 0.46^a^ | 3.43 ± 0.40 |
| On litter (%) | 77.70 ± 0.87^ab^ | 78.19 ± 0.74^a^ | 74.51 ± 0.79^b^ | 77.02 ± 0.76^ab^ | 72.04 ± 1.05 |
| **Day 42** |  |  |  |  |  |
| All enrichments (%) | 22.65 ± 1.17^a^ | 24.81 ± 1.15^ab^ | 29.21 ± 1.16^b^ | 26.35 ± 1.15^ab^ | 31.29 ± 5.14 |
| On platform (%) | 9.11 ± 0.77 | 9.19 ± 0.76 | 11.08 ± 0.77 | 8.57 ± 0.76 | 14.12 ± 2.32 |
| Under platform and ramp (%) | 6.22 ± 1.00^a^ | 10.36 ± 0.98^ab^ | 9.03 ± 0.99^ab^ | 12.04 ± 0.98^b^ | 6.58 ± 1.08 |
| On scale (%) | 5.25 ± 0.48^ab^ | 4.72 ± 0.48^a^ | 6.95 ± 0.48^b^ | 4.49 ± 0.48^a^ | 2.83 ± 0.47 |
| On litter (%) | 83.38 ± 0.82^ab^ | 85.41 ± 0.81^a^ | 80.03 ± 0.81^b^ | 85.84 ± 0.81^a^ | 75.29 ± 12.38 |
| **Day 56** |  |  |  |  |  |
| All enrichments (%) | 23.48 ± 1.21^a^ | 24.07 ± 0.91^a^ | 28.97 ± 0.93^b^ | 25.96 ± 0.92 | 28.93 ± 8.03 |
| On platform (%) | 8.25 ± 0.57^ab^ | 7.58 ± 0.44^a^ | 10.02 ± 0.45^b^ | 8.42 ± 0.44^ab^ | 12.32 ± 3.43 |
| Under platform and ramp (%) | 5.56 ± 0.76^a^ | 10.53 ± 0.57^b^ | 11.47 ± 0.58^b^ | 12.09 ± 0.57^b^ | 8.23 ± 2.28 |
| On scale (%) | 6.91 ± 0.66^a^ | 5.03 ± 0.50^ab^ | 6.09 ± 0.51^a^ | 3.76 ± 0.49^b^ | 2.86 ± 0.79 |
| On litter (%) | 80.27 ± 1.22^a^ | 86.27 ± 0.80^b^ | 82.51 ± 0.82^ac^ | 86.07 ± 0.80^b^ | 79.27 ± 21.99 |

^1^ Raw means are presented; due to a low sample size, strain T (ADG_0-61_ =19.78 g/day) was not included in the statistical models.

^a ≠ b^ within rows at adjusted P < 0.05

**Supplementary Table 9.** Enrichment use, assessed as the percentage of birds in the pen using each enrichment (back-transformed LS-means ± SEM), within CONV strains according to target weight.

|  | **A^1^** | **B** | **C** |
| --- | --- | --- | --- |
| **Target weight 1** |  |  |  |
| All enrichments (%) | 22.43 ± 3.55 | 23.11 ± 0.80 | 23.06 ± 0.88 |
| On platform (%) | 8.96 ± 1.42 | 9.20 ± 0.42^a^ | 7.25 ± 0.46^b^ |
| Under platform and ramp (%) | 7.45 ± 1.18 | 6.63 ± 0.49 | 8.28 ± 0.54 |
| On scale (%) | 3.69 ± 0.06 | 5.72 ± 0.41 | 5.26 ± 0.44 |
| On litter (%) | 85.03 ± 13.44 | 83.50 ± 0.65 | 85.04 ± 0.70 |
| **Target weight 2** |  |  |  |
| All enrichments (%) | 21.41 ± 4.56 | 19.67 ± 1.09 | 20.13 ± 1.06 |
| On platform (%) | 6.82 ± 1.45 | 7.63 ± 0.56 | 6.95 ± 0.55 |
| Under platform and ramp (%) | 7.54 ± 1.61 | 6.74 ± 0.68 | 8.82 ± 0.66 |
| On scale (%) | 5.62 ± 1.20 | 4.59 ± 0.52 | 4.34 ± 0.51 |
| On litter (%) | 85.92± 18.32 | 87.35 ± 0.91 | 88.32 ± 0.83 |

^1^ Raw means are presented; due to a low sample size, strain A (ADG_0-47_ = 62.65 g/day) was not included in the statistical models.

^a ≠ b^ within rows at adjusted P < 0.05

**Supplementary Table 10.** Enrichment use, assessed as the percentage of birds in the pen using each enrichment (back-transformed LS-means ± SEM), within FAST strains according to target weight.

|  | **F** | **G** | **I** | **M** |
| --- | --- | --- | --- | --- |
| **Target weight 1** |  |  |  |  |
| All enrichments (%) | 20.23 ± 0.91^a^ | 24.19 ± 0.89^b^ | 25.52 ± 0.89^b^ | 22.68 ± 0.95^ab^ |
| On platform (%) | 5.73 ± 0.47^a^ | 9.07 ± 0.46^b^ | 9.37 ± 0.46^b^ | 9.49 ± 0.49^b^ |
| Under platform and ramp (%) | 8.93 ± 0.56^a^ | 8.40 ± 0.54^a^ | 8.39 ± 0.54^a^ | 4.73 ± 0.59^b^ |
| On scale (%) | 4.71 ± 0.45^a^ | 5.55 ± 0.45^ab^ | 6.12 ± 0.45^b^ | 4.97 ± 0.47 |
| On litter (%) | 88.60 ± 0.73^a^ | 84.17 ± 0.71^b^ | 82.83 ± 0.71^b^ | 82.97 ± 0.76^b^ |
| **Target weight 2** |  |  |  |  |
| All enrichments (%) | 18.14 ± 1.17^ac^ | 22.96 ± 1.13^bc^ | 25.13 ± 1.15^b^ | 18.85 ± 1.51^c^ |
| On platform (%) | 6.04 ± 0.61^a^ | 9.46 ± 0.59^b^ | 8.46 ± 0.60^b^ | 8.78 ± 0.78^ab^ |
| Under platform and ramp (%) | 7.95 ± 0.73^ab^ | 8.49 ± 0.70^a^ | 8.58 ± 0.72^a^ | 4.26 ± 0.96^b^ |
| On scale (%) | 3.57 ± 0.56^a^ | 4.22 ± 0.54^a^ | 6.30 ± 0.56^b^ | 3.46 ± 0.70^a^ |
| On litter (%) | 89.80 ± 0.91^a^ | 85.46 ± 0.88^b^ | 83.35 ± 0.90^b^ | 85.50 ± 1.25^ab^ |

^a ≠ b ≠ c^ within rows at adjusted P < 0.05

**Supplementary Table 11.** Enrichment use, assessed as the percentage of birds in the pen using each enrichment (back-transformed LS-means ± SEM), within MOD strains according to target weight.

|  | **E** | **H** | **O** | **S** |
| --- | --- | --- | --- | --- |
| **Target weight 1** |  |  |  |  |
| All enrichments (%) | 20.91 ± 0.90^a^ | 23.47 ± 0.87^ab^ | 25.01 ± 0.89^b^ | 23.25 ± 0.90^ab^ |
| On platform (%) | 8.24 ± 0.47 | 9.03 ± 0.45 | 8.86 ± 0.46 | 8.77 ± 0.47 |
| Under platform and ramp (%) | 6.34 ± 0.55 | 7.29 ± 0.53 | 8.25 ± 0.54 | 7.63 ± 0.55 |
| On scale (%) | 4.83 ± 0.45^a^ | 5.43 ± 0.44^ab^ | 6.53 ± 0.45^b^ | 6.07 ± 0.45^ab^ |
| On litter (%) | 85.46 ± 0.72 | 83.73 ± 0.70 | 83.22 ± 0.71 | 84.32 ± 0.72 |
| **Target weight 2** |  |  |  |  |
| All enrichments (%) | 20.10 ± 1.17^a^ | 22.63 ± 1.26^ab^ | 25.13 ± 1.14^b^ | 22.98 ± 1.18^ab^ |
| On platform (%) | 7.83 ± 0.61 | 9.07 ± 0.65 | 9.33 ± 0.58 | 9.11 ± 0.58 |
| Under platform and ramp (%) | 6.51 ± 0.74 | 6.70 ± 0.79 | 8.31 ± 0.69 | 8.31 ± 0.69 |
| On scale (%) | 4.15 ± 0.56^a^ | 5.75 ± 0.59^ab^ | 6.35 ± 0.53^b^ | 5.14 ± 0.54^ab^ |
| On litter (%) | 86.69 ± 0.97 | 84.04 ± 1.07 | 83.11 ± 0.87 | 85.23 ± 0.87 |

^a ≠ b^ within rows at adjusted P < 0.05

**Supplementary Table 12.** Enrichment use, assessed as the percentage of birds in the pen using each enrichment (back-transformed LS-means ± SEM), assessed as the percentage of birds in the pen using each resource, within SLOW strains according to target weight.

|  | **D** | **J** | **K** | **N** | **T^1^** |
| --- | --- | --- | --- | --- | --- |
| **Target weight 1** |  |  |  |  |  |
| All enrichments (%) | 21.60 ± 0.86^a^ | 24.65 ± 0.89^ab^ | 27.13 ± 0.89^b^ | 25.98 ± 0.88^b^ | 36.70 ± 7.06 |
| On platform (%) | 9.04 ± 0.45 | 9.74 ± 0.46 | 10.40 ± 0.46 | 8.94 ± 0.46 | 19.25 ± 3.71 |
| Under platform and ramp (%) | 5.12 ± 0.53^a^ | 9.30 ± 0.55^bc^ | 8.31 ± 0.55^b^ | 10.70 ± 0.54^c^ | 6.35 ± 1.22 |
| On scale (%) | 5.28 ± 0.44^a^ | 4.84 ± 0.45^a^ | 6.85 ± 0.45^b^ | 4.62 ± 0.45^a^ | 2.46 ± 0.05 |
| On litter (%) | 83.44 ± 0.70^ab^ | 84.60 ± 0.72^a^ | 81.14 ± 0.71^b^ | 84.66 ± 0.71^a^ | 68.37 ± 2.25 |
| **Target weight 2** |  |  |  |  |  |
| All enrichments (%) | 24.36 ± 1.26^ab^ | 22.35 ± 1.12^a^ | 27.47 ± 1.11^b^ | 24.27 ± 1.12^ab^ | 35.43 ± 8.13 |
| On platform (%) | 9.92 ± 0.65 | 7.95 ± 0.58 | 9.94 ± 0.58 | 8.70 ± 0.58 | 18.78 ± 4.31 |
| Under platform and ramp (%) | 4.29 ± 0.78^a^ | 8.66 ± 0.70^b^ | 9.70 ± 0.69^b^ | 10.24 ± 0.69^b^ | 8.78 ± 2.01 |
| On scale (%) | 7.10 ± 0.59^a^ | 5.30 ± 0.54^ab^ | 6.76 ± 0.53^a^ | 4.04 ± 0.54^b^ | 2.35 ± 0.05 |
| On litter (%) | 78.09 ± 1.15^a^ | 86.23 ± 0.88^b^ | 82.13 ± 0.^87ac^ | 85.92 ± 0.87^b^ | 78.21 ± 2.40 |

^1^ Raw means are presented; due to a low sample size, strain T (ADG_0-61_ =19.78 g/day) was not included in the statistical models.

^a ≠ b ≠ c^ within rows at adjusted P < 0.05
